# Frequent horizontal chromosome transfer between asexual fungal insect pathogens

**DOI:** 10.1101/2023.09.18.558174

**Authors:** Michael Habig, Anna V. Grasse, Judith Müller, Eva H. Stukenbrock, Hanna Leitner, Sylvia Cremer

**Author notes:** Corresponding authors: Michael Habig,; Sylvia Cremer.

## Abstract

Entire chromosomes are typically only transmitted vertically from one generation to the next. The horizontal transfer of such chromosomes has long been considered improbable, yet gained recent support in several pathogenic fungi where it may affect the fitness or host specificity. To date, it is unknown how these transfers occur, how common they are and whether they can occur between different species. In this study, we show multiple independent instances of horizontal transfers of the same accessory chromosome between two distinct strains of the asexual entomopathogenic fungus *Metarhizium robertsii* during experimental co-infection of its insect host, the Argentine ant. Notably, only the one chromosome – but no other – was transferred from the donor to the recipient strain. The recipient strain, now harboring the accessory chromosome, exhibited a competitive advantage under certain host conditions. By phylogenetic analysis we further demonstrate that the same accessory chromosome was horizontally transferred in a natural environment between *M. robertsii* and another congeneric insect pathogen, *M. guizhouense*. Hence horizontal chromosome transfer is not limited to the observed frequent events within species during experimental infections but also occurs naturally across species. The transferred accessory chromosome contains genes that might be involved in its preferential horizontal transfer, encoding putative histones and histone-modifying enzymes, but also putative virulence factors that may support its establishment. Our study reveals that both intra- and interspecies horizontal transfer of entire chromosomes is more frequent than previously assumed, likely representing a not uncommon mechanism for gene exchange.

**Significance Statement:** The enormous success of bacterial pathogens has been attributed to their ability to exchange genetic material between one another. Similarly, in eukaryotes, horizontal transfer of genetic material allowed the spread of virulence factors across species. The horizontal transfer of whole chromosomes could be an important pathway for such exchange of genetic material, but little is known about the origin of transferable chromosomes and how frequently they are exchanged. Here, we show that the transfer of accessory chromosomes - chromosomes that are non-essential but may provide fitness benefits - is common during fungal co-infections and is even possible between distant pathogenic species, highlighting the importance of horizontal gene transfer via chromosome transfer also for the evolution and function of eukaryotic pathogens.

## Introduction

Eukaryotic genomes consist of essential core genomes and, in some species, may also contain accessory chromosomes that are not essential. Accessory chromosomes are defined by their presence/absence polymorphism within a species and are widely observed in fungi, plants, and animals. These unique chromosomes (also known as supernumerary, lineage-specific, conditionally dispensable or B chromosomes) can provide additional functions such as virulence factors and are particularly common in fungal plant pathogens (1–4). The fitness effects of fungal accessory chromosomes can vary from negative to positive and appear to depend on the plant host species (5–9). Many fungal pathogens that harbor accessory chromosomes can infect a range of host species, so the presence/absence polymorphism of the accessory chromosomes is currently thought to be mainly due to their varying fitness effects in different plant host species. Despite their importance, the origin of accessory chromosomes is unknown; they may have originated from within the own genome (2) or may have been horizontally acquired from another species. The latter is supported by genomic comparisons for a few species (8, 10). Experimentally, however, horizontal transfer of accessory chromosomes has only been observed in very few fungal species. In all these cases, the transfer was observed during growth *in vitro* and at very low frequencies, requiring the use of inserted selectable fungicide-resistance markers in these studies (10–13). Hence, to date horizontal transfer of an entire chromosome has not been observed during the natural stages of the life cycle of a fungus, e.g., during infections of a host for a pathogenic fungus. As a result, neither the frequency nor the underlying mechanisms of horizontal chromosome transfer between fungal pathogens are well-understood.

Horizontal transfer of genetic material between different species was first recognized among bacteria, but has since then been found to occur also in the genomes of many eukaryotes (14–17). Transferred genetic material can have an adaptive advantage (18–21), such as the ToxA gene, which encodes an effector that interferes with the host plant’s immune system, and was horizontally transferred between three fungal pathogens of wheat (22, 23). The mechanisms by which such horizontal transfer of genetic material can occur in eukaryotes are less well established. In general, the horizontal transfer of genetic material between different species of eukaryotes is believed to be mostly the result of hybridization by sexual processes. The highly orchestrated nature of sexual processes may however limit which species can produce hybrids with each other and therefore the frequency and direction of horizontal transfers. Parasexuality in fungi, on the other hand, represents an additional process for horizontal transfer that would presumably allow for greater compatibility. Here, vegetative cells fuse, eventually resulting in cells with multiple nuclei that either exchange genetic material (24) or fuse (karyogamy) (25, 26). Prominent examples of such parasexual cycles are the human pathogen *Candida albicans* and *Aspergillus fumigatus* (18, 27–29). Here, the fusion of the nuclei might be followed by mitotic recombination and random chromosome losses that will eventually re-establish a stable ploidy level (18, 19, 30). Therefore, the horizontal transfer of genetic material in eukaryotes is important, but observing such transfer events is very difficult because they seem to occur very rarely.

Here, we ask whether, how frequent and to what extent genetic material can be horizontally transferred between asexual lineages of a fungal pathogen during co-infections of their insect host. To this end, we studied common insect pathogens of the genus *Metarhizium* (Ascomycota), which frequently infect and kill insects and are often used as biocontrol agents against insect pests (31). In addition to their parasitic lifestyle in insects, some are also associated with plants as rhizosphere colonizers and root endophytes (31). Here we used a set of fungal strains derived from a selection experiment, in which Argentine ants (*Linepithema humile*) were co-infected with a mixture of six strains of *M. robertsii* and *M. brunneum* and then kept under conditions either allowing only the ant’s individual immune defenses to act, or both their individual and cooperative social defenses (32). By analyzing the ancestral and the evolved fungal strains resulting from that experiment we show that i) horizontal transfer of an accessory chromosome occurred frequently between two distinct strains of *M. robertsii*, and ii) only the accessory chromosome was transferred and spread through the population over ten host infection cycles depending on the presence or absence of the ant’s social immune defenses. iii) Lastly, a phylogenetic analysis of 36 strains across the genus *Metarhizium* revealed that the same accessory chromosome, which easily transmits within species of *M. robertsii*, was horizontally transferred to a distant, congeneric *M. guizhouense*. Taken together, our results indicate that an accessory chromosome in *Metarhizium* is highly mobile and encodes factors that may influence its mobility as well as putative virulence factors.

## Results

In a previous experiment we performed twenty serial co-infections (passages) of Argentine ants with six different strains of *M. robertsii* and *M. brunneum* in two treatment groups, consisting of either one individual ant (treatment: individual) or one ant in the presence of two nestmates performing social grooming (treatment: social), with ten replicates for each treatment (see Fig 1A for a depiction of the experimental procedure) (Stock et al. (31)). As a result, we found that a co-infecting mix of *M. robertsii* and *M. brunneum* shows different phenotypic adaptations to the individual versus the social immune defenses of their Argentine ant hosts (31). We here performed a detailed molecular analysis of all six ancestral and the 24 resulting evolved fungal strains to determine the genetic changes associated with this selection experiment.

**Fig. 1:**
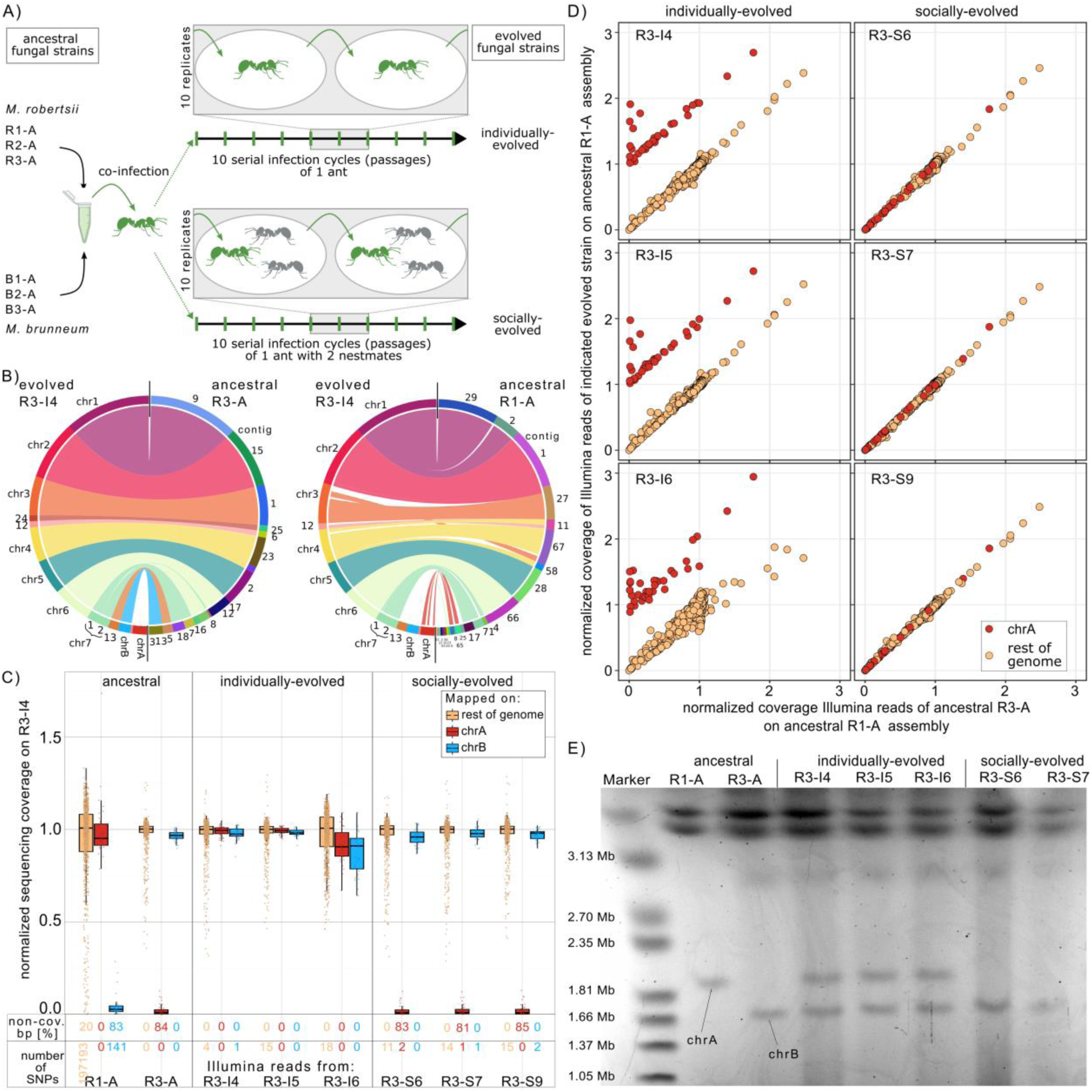
Multiple independent transfers of accessory chrA from the ancestral *M. robertsii* R1-A to evolved *M. robertsii* R3 during an experimental co-infection of an ant host. A) Experimental procedure of the selection experiment performed by Stock et al. (32). Argentine ants (green) were exposed to a mix of six strains – three *M. robertsii* and three *M. brunneum* – and kept either alone (individual treatment, n=10 replicate lines) or with two nestmates (grey; social treatment, n=10 replicate lines). The produced infectious spores were used to expose ants in ten serial infection cycles as described in (32). We here performed whole genome sequencing of the six ancestral and the individually- and socially-evolved strains at the end of the experiment. B) Synteny blot of the nanopore-based assemblies of the evolved R3-I4 compared to the two ancestral R1-A and R3-A. Accessory chrA is missing in R3-A but shows synteny to contigs in R1-A. C) Normalized Illumina sequencing coverage mapped on the evolved R3-I4 nanopore-based assembly in 50 kb windows for chrA, chrB, and the rest of the genome. The fraction of genomic compartments lacking a single sequencing read (non-cov. bp) and the number of SNPs/InDels absent in the ancestral R3-A strain are given. D) Normalized sequencing coverage comparison of Illumina reads mapped on the ancestral R1-A nanopore-based assembly in 50 kb windows. Only 50 kb windows that are syntenic to chrA of R3-IA exhibited changes in sequencing coverage in the three individually-evolved R3 strains, indicating that no other genetic material was transferred to them from the ancestral R1-A. Note: The plots excluded the rDNA cluster due to its high coverage for clarity. E) Pulsed-field gel electrophoresis (PFGE) image of chromosomes from the ancestral R1-A and R3-A, as well as all three individually-evolved and two socially-evolved R3 strains. All three independent individually-evolved R3 strains contained both chrA and chrB.

### Frequent horizontal transfer of solely accessory chromosome A in *M. robertsii* during insect infection

Preliminary analysis on fragmented Illumina-based assemblies for *M. robertsii* and *M. brunneum* indicated that horizontal transfer may have occurred between two strains of *M. robertsii* (R1 and R3), but not between the other co-infecting strains. In order to further analyze this potential transfer, we generated nanopore-based assemblies for two ancestral strains (R1-A and R3-A) and one evolved strain (R3 from individual treatment replicate 4, i.e. R3-I4). These near-chromosome-level assemblies (Fig. S1 A-C, Table S1) showed a high degree of synteny with the only published chromosome-level assembly for a *Metarhizium* species, the related *M. brunneum* (Fig. S1 D-F) (33), confirming our assembly procedure. We obtained the best assembly, with the lowest number of contigs, for the evolved R3-I4 strain, and consequently, renamed our R3-I4 main contigs based on their synteny with *M. brunneum*’s chromosome-level assemblies. Interestingly one complete chromosome with telomeres at both ends (1.81 Mb) was absent in the ancestral R3-A strain but showed synteny with eleven contigs of the ancestral R1-A (Fig. 1B). Hence this chromosome showed presence/absence polymorphism in *M. robertsii* and is therefore accessory, and we termed this accessory chromosome A (chrA). Based on the synteny there were no single nucleotide polymorphism (SNPs) on chrA between R1-A and R3-I4 (Fig. S2 A-C). Remarkably, the absence of SNPs for accessory chrA contrasted with the high SNP density observed in all other syntenic chromosomes between R1-A and R3-I4. Additionally, we found another accessory chromosome (1.64 Mb) in the evolved R3-I4, present in the R3-A ancestral strain but absent in the R1-A ancestral strain (Fig 1. B). We termed this chromosome accessory chromosome B (chrB).

To confirm that chrA is indeed non-existent in the ancestral R3-A, and likewise chrB in R1-A – that is, their absence in the assembled sequences not reflecting an assembly error – we mapped Illumina sequencing reads to the R3-I4 assembly, which contains both accessory chrA and chrB (Figure 1 C). Both chrA and chrB showed minimal sequencing coverage when mapped with the reads from the ancestral R3-A and R1-A, respectively, with the majority of chrA and chrB showing no sequencing read (84% and 83% of bases without a single sequencing read, respectively) and low median normalized coverage in 50-kb windows (0.9% and 2.6%, respectively) (Fig. 1 C). Therefore, we concluded that the accessory chromosomes chrA and chrB were indeed absent in the ancestral R3-A and R1-A ancestral strains, respectively. Thus, the two ancestral strains possess different accessory chromosomes, with chrA having been horizontally transferred from the R1-A ancestral strain to the R3 strain, resulting in the evolved R3-I4.

Sequencing coverage analysis revealed that all three R3 strains that had independently evolved under the individual treatment (R3-I4, R3-I5, R3-I6) acquired the accessory chrA from the ancestral R1-A strain, while chrA was not present at the end of the experiment in any of the three R3 strains that independently evolved under the social treatment (R3-S6, R3-S7, R3-S9; Fig 1 C), a pattern that was further confirmed by Pulsed-field Gel Electrophoresis (PFGE; Fig. 1 E). While this clearly shows the transfer of chrA, it does not yet exclude the possibility that other genetic material could have been transferred at the same time. To test for this, we, first, compared the normalized sequencing coverage of 50 kb windows between the six evolved R3 strains with their R3-A ancestor strain, when mapped to the ancestral R1-A assembly (Fig. 1 D). We observed an increase in sequencing coverage by approx. one normalized sequencing coverage for those sequences syntenic to accessory chrA in all three chrA-recipient R3 strains. This suggests that only chrA was transferred and that the recipient strains now contained a single copy of chrA. Genomic regions of R1-A that were not syntenic to chrA showed no change in coverage between the ancestral and the evolved R3, indicating the absence of additional transferred genetic material. Second, we analyzed the presence of R1-A-specific SNPs in the individually-evolved R3 strains. Despite the high number of 194,334 SNPs differentiating the ancestral strains R1-A and R3-A (Table S2), we did not identify any R1-A-specific SNPs in the evolved R3 strains, ruling out chromosome replacement or mitotic recombination events (Fig. S3). Hence solely chrA was horizontally transferred independently three times while no other genetic material was transferred from R1-A to the individually-evolved R3 strains.

### The methylation pattern of the chrA is partially retained in the recipient strain

The intraspecific horizontal transfer of chrA from R1-A to R3 during our experiment represents the transfer of an entire chromosome from one genomic background to another. Here we have the unique opportunity to study how epigenetic marks were affected by such a horizontal transfer, because we have access to both ancestral strains as well as the recipient strain that now harbors chrA. We focused on DNA-methylation, which is known to vary between different *Metarhizium* strains (34–36). We hypothesized that chrA may possess a distinct DNA methylation pattern compared to the rest of the genome in the donor strain and that its pattern might be affected by the horizontal transfer to R3. Using Nanopore sequencing data, we identified cytosine methylation in the CpG context. Overall, the ancestral R1-A strain exhibited significantly higher methylation levels than the ancestral R3-A strain (Fisher exact test, p<4.6 × 10^−15^) (Fig S4 A, Table S3). Furthermore, in both the ancestral R1-A strain and the evolved R3-I4 strain chrA showed significantly higher methylation levels compared to the rest of the genome (Fisher exact test, p<4.6 × 10^−15^ for both comparisons). Upon closer examination, we found that the subtelomeric regions of chrA in R3-I4 retained similar levels of methylation compared to R1-A, while the central portions displayed methylation levels similar to the genome-wide average (Fig S4 B). Although the methylation level on chrA was substantially lower in the evolved R3-I4 strain compared to the ancestral R1-A, the majority (549 out of 741) of methylated sites in R3-I4 were also methylated in R1-A (Fig. S4 C). These results led us to conclude that the transferred chrA in the recipient R3-I4 strain retained some of the methylation pattern of its donor, the ancestral R1-A strain, although the absolute level of methylation decreased upon transfer to the recipient strains. This is therefore an example of how the genomic context can influence epigenetic marks, potentially also affecting gene expression.

### chrA provides a competitive advantage in individual ant hosts

Next, we wanted to understand the temporal dynamics of the horizontal transfers and whether recipient strains had a competitive advantage or disadvantage compared to the same strain without chrA. To this end, we performed a detailed analysis of the clone diversity at passages 1, 3, 5 and 10 testing the proportion of R3 spores with present versus absent chrA for the six above-mentioned R3-strains that had persisted until passage 10, plus another two that had persisted at least until passage 5, but had become outcompeted by passage 10 (R3-S1, R3-S8), see Stock et al. (32). We found that chrA was transferred from R1 to R3 in five out of the eight selection lines (in all three individually-evolved, as well as in two of five of the socially-evolved) (Fig 2 A-B). Hence, horizontal transfer occurred frequently. The main difference between the individual and social immunity treatments arose, however, in the persistence of the R3 strains that had acquired the chrA. When adapting to individual ants, chrA spread very quickly through the fungal population. After ten passages, chrA was present in all R3 spores across all three replicates. This contrasts with the lack of the spread of chrA in R3 strains adapting to social ants. Despite being transferred from R1 to R3 in two of the five replicates (R3-S1, R3-S6), R3 became completely extinct in the socially-evolved replicate S1, whilst in replicate S6, R3 had outcompeted all other strains at the end of the experiment, but the winning R3 spores were the ones not containing chrA. This means that acquiring chrA improved the competitive ability of the evolved R3 against the R3 without chrA in individual ant hosts, yet when the exposed ants were with caretaking nestmates, the opposite was the case. This indicates that social immunity did not interfere with chrA transfer rates, but with its establishment and long-term persistence in the pathogen population, providing a further example of the modulatory power of social immunity on the competitive ability of its coinfecting pathogens (37).

**Fig. 2:**
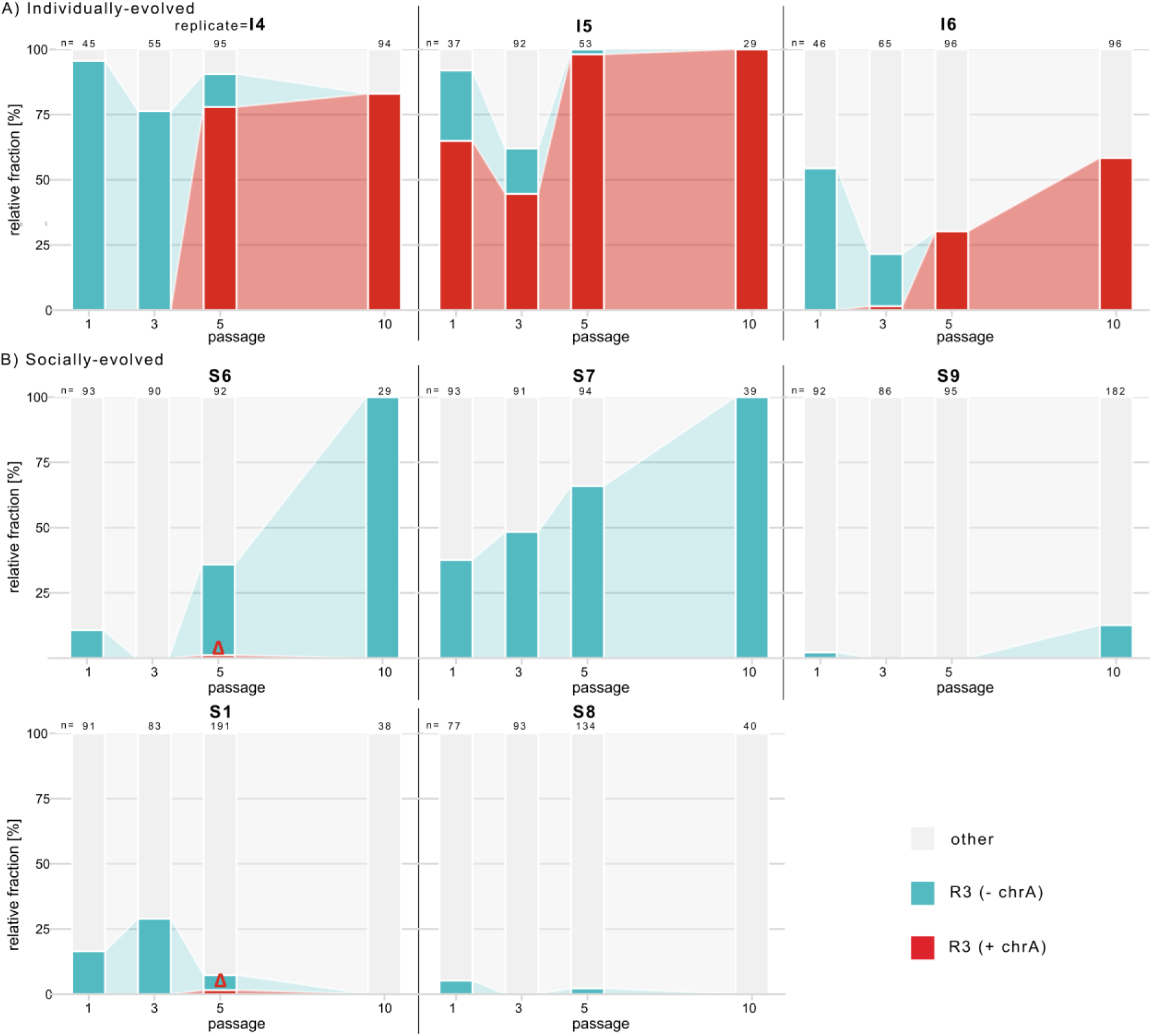
Comparative dynamics of R3 spores with either chrA present or absent over the course of the selection experiment. Proportion of R3 spores with integrated (red) or absent (turquoise) chrA in comparison to the remaining strains (grey) analyzed in passages 1, 3, 5, and 10 in (A) individual ant hosts (I4, I5, I6) or (B) ants with nestmates providing sanitary care (upper row: replicate lines, in which R3 persisted until passage 10 (S6, S7, S9); lower row: replicate lines, in which R3 still persisted in passage 5, but was outcompeted until passage 10 (S1, S8)). The value of n represents the total number of genotyped single-spore derived colonies (n=total of 2626 spores). Please note that the occurrence of chrA in R3 in two of the socially-evolved replicate lines, where it later failed to establish, is highlighted by a red triangle.

### ChrA shows presence/absence polymorphism in the genus *Metarhizium*

Our experimental data therefore revealed that chrA is easily horizontally transferred between different *M. robertsii* strains during co-infection in an insect host. We hypothesized that the accessory chromosome in *M. robertsii* might also be transferable between different species, since many species of *Metarhizium* have a broad host range (31), making congeneric co-infections a likely common scenario in the field. We therefore tested if we find evidence for chrA being present across the genus *Metarhizium*, by use of phylogenetic analysis of the six ancestral strains of our experiment and all 30 published sequences of multiple *Metarhizium* strains and species (*M. robertsii*, *M. anisopliae*, *M. brunneum*, *M. guizhouense*, *M. majus*, *M. acridum*, and *M. album*), all isolated from natural populations from the field (Table S4).

Available assemblies and short sequencing reads were mapped onto the assembly of our individually-evolved *M. robertsii* strain R3-I4 that contains both, chrA and chrB. For each strain, we determined the coverage as a fraction of bases covered (excluding transposable elements) for both, chrA (Fig. 3A) and chrB (Fig. S5A). Within *M. robertsii*, chrA shows a presence/absence polymorphism, with 100% coverage and an absence of SNPs in our ancestral R1-A donor strain and with very low coverage in the ancestral R3-A, indicating its absence. Among other *M. robertsii* strains, the relative coverage of chrA varied from 39.9%. to 70.3%. Similarly, in *M. anisopliae*, the relative coverage ranged from 12.7% to 51.5%, while in *M. brunneum*, it ranged from 2.8% to 61%. In *M. majus*, chrA had coverage of 47.3% of all bases, whereas it was poorly covered in *M. acridum* and *M. album* (1.4-1.9%). Interpreting the intermediary coverage is difficult due to availability of only very fragmented assemblies for these strains. Hence it is unclear whether the sequences are located on one chromosome (which would then be accessory) or distributed within the genome.

**Fig. 3:**
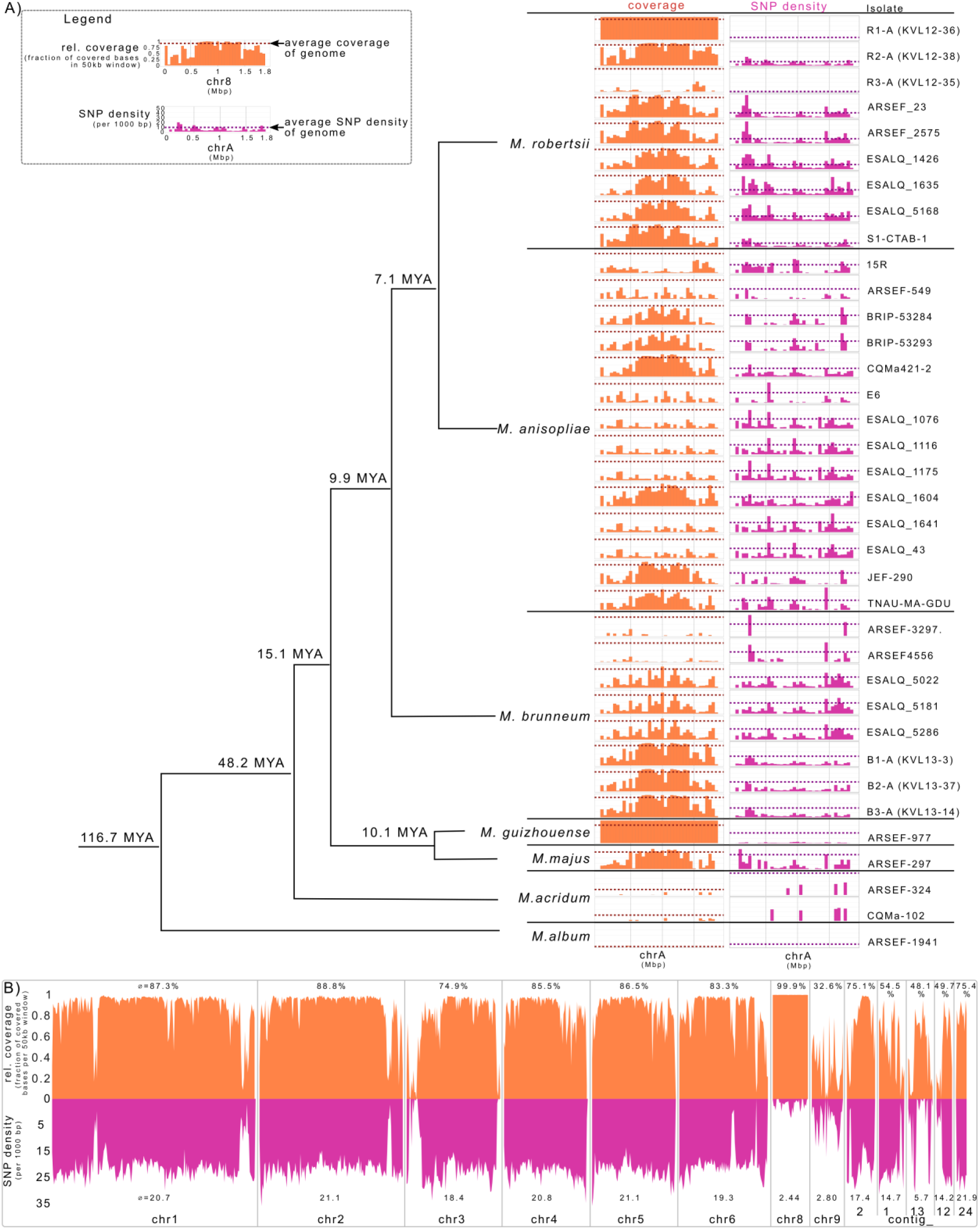
Distribution of chrA sequences across *Metarhizium* species. A) Phylogeny of published *Metarhizium* strains (from Hu *et al*. (38)), for which assemblies or sequencing reads were available. These assemblies or sequencing reads were mapped onto the near-chromosome level assembly of *M. robertsii* R3-I4, which contains both accessory chrA and chrB. The fraction of covered bases of chrA (orange) and the density of SNPs (pink) were determined in 50 kb windows. Dotted lines represent respective genome-wide averages of the covered sequences and the SNP density. Transposable elements (TEs) were excluded from the analysis. MYA: Million years ago. B) Relative sequence coverage of Illumina reads from *M. guizhouense* ARSEF-977 mapped to the evolved *M. robertsii* R3-I4 strain assembly. The fraction of bases covered per 50 kb window is shown in orange while SNP density per 1000 bp in 50 kb windows is shown in pink. ChrA exhibits higher average sequence coverage and lower SNP density in *M. guizhouense* compared to the rest of the genome (see also Fig S5 A for chrB).

Interestingly, *M. guizhouense* exhibited complete coverage (99.9%) of all bases on chrA, as well as a very low number of SNPs. This clearly demonstrates that chrA is shared between *M. robertsii* and *M. guizhouense*, despite their separation 15.1 Mio years ago. Performing the same analysis on chrB (Fig. S5A) could not identify the presence of this second accessory chromosome of *M. robertsii* in any other of the 35 strains than in our ancestral R3-A strain. We could also not detect any relation with the presence/absence of the two accessory chromosomes. In conclusion, there is a high variation in the presence of sequences of the accessory chrA and chrB that does not relate to the phylogeny and therefore might be indicative of either frequent losses of these chromosomes or horizontal transfer.

### Horizontal transfer of chrA across *Metarhizium* species borders

Accessory chrA showed a markedly reduced SNP density between *M. robertsii* and *M. guizhouense* compared to the rest of the genomes (2.44 compared to 19.2 SNPs per 1000 bp, respectively) (Fig. 3 B, Table S4), which could be indicative of a horizontal transfer. We therefore analyzed chrA in *M. guizhouense* in more detail. Mapping the short sequencing reads of *M. guizhouense* revealed that chrA displayed approximately double the normalized sequencing coverage compared to the rest of the genome (Fig. 4 A). These observations strongly suggest that the sequences present on chrA are duplicated in *M. guizhouense*, indicating the presence of a disomic state for chrA in this strain. PFGE confirmed the presence of at least two small chromosomes similar in size to the accessory chromosomes chrA and chrB in *M. guizhouense* (Fig S5 B). Therefore, we conclude that chrA exists in two copies in *M. guizhouense*, and the relatively low SNP density separating chrA in *M. robertsii* from *M. guizhouense*, compared to the significantly higher SNP density in the rest of the genome, likely reflects a shorter separation time from their respective syntenic sequences for chrA than for the rest of the genome.

**Fig. 4:**
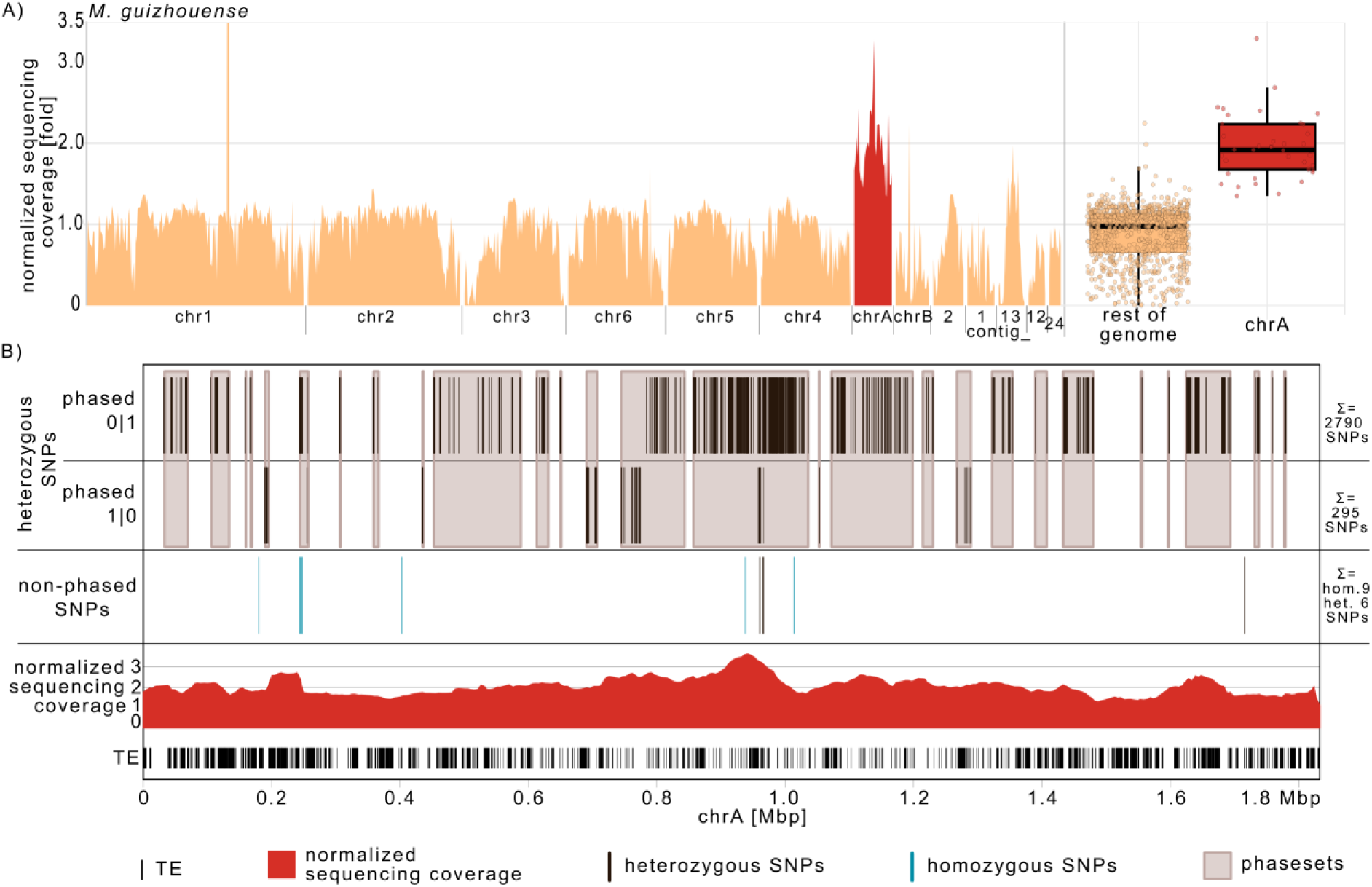
Sequencing coverage and SNP density indicate two copies of chrA in *M. guizhouense* that differ from each other in their SNP density. A) Normalized Illumina sequencing coverage of *M. guizhouense* ARSEF-977 on the *M. robertsii* R3-I4 assembly, presented separately for each contig and summarized for chrA and the rest of the genome in 50 kb windows. The coverage of the rDNA cluster is not displayed in full, for visual clarity. B) Phasing information of *M. guizhouense* SNPs located outside of transposable elements (TEs) on the *M. robertsii* R3-I4, shown in individual phasesets. Most phasesets exclusively contain SNPs from one phase. The figure also depicts the normalized sequencing coverage in 50 kb (5 kb sliding) windows, along with the location of transposable elements (hom = homozygous, het = heterozygous).

Next, we wondered if the two copies of chrA in *M. guizhouense* differ in respect to their separation time from chrA in *M. robertsii* (R1-A) by determining their SNP density separately for each copy. We therefore phased the heterozygous SNPs of *M. guizhouense*, i.e. we tried to determine on which of the two copies of chrA a SNP is located. Phasing SNPs is most efficient using long sequencing reads and we therefore used long PacBio reads. The result of phasing are separate phasesets, for which the location, i.e. on which of the two chromosome copies, of the SNPs relative to the other SNPs of the same phaseset is known. Among the total of 3090 heterozygous SNPs on chrA, 2790 were phased as 0|1 and 294 as 1|0. Interestingly, the vast majority of phasesets contained SNPs of only one phase, not both (25 out of 28 phasesets in total). Merely three phasesets contained SNPs of both phases, i.e. SNPs that for which the location is on the two different copies. The largest phaseset with SNPs of both phases (chrA: 587009-1035720) consisted of 1076 phased SNPs, with 1064 (98.9%) phased as 0|1 and only 12 (1.1%) phased as 1|0 (Fig. 4 B). Such an imbalanced distribution of SNPs between the two copies of chrA in one phaseset would be highly unlikely if both copies of chrA had the same density and distribution of SNPs (p < 2.2×10^−16^, binomial test). Consequently, we concluded that the two copies of chrA differ from each other in the density and distribution of SNPs. Because we had several phasesets along chrA, we could not directly assign the phases 0|1 or 1|0 each to one copy of chrA. We therefore utilized the number of mixed phasesets (containing SNPs from both phases) and unmixed phasesets (containing SNPs from only one phase) to estimate the number of SNPs on each of the two copies of chrA. Out of 28 phasesets, 25 (89.3%) exclusively contained SNPs from the same phase, while three (10.7%) contained SNPs from both phases. Therefore, one copy (copy a) harbours an estimated 2769 SNPs (calculated as 3084 phased SNPs × 25/28 + 6 non-phased SNPs + 9 homozygous SNPs), whereas the other copy (copy b) contains approximately 345 SNPs (calculated as 3084 phased SNPs × 3/28 + 6 non-phased SNPs + 9 homozygous SNPs). Therefore, the two copies of chrA in *M. guizhouense* differ in their SNP density from each other as well as from the rest of the genome compared to *M. robertsii*, indicating different separation times for the two copies of chrA.

When did the two copies of chrA in *M. guizhouense* separate from the chrA found in *M. robertsii*? The estimated divergence between *M. guizhouense* and *M. robertsii* is 15.1 million years ago (38), which allows us to calibrate the molecular clock based on the rest of the genome for a comparison between *M. robertsii* and *M. guizhouense*. Using this calibration, we estimated that the separation of the two copies of chrA in *M. guizhouense* from chrA in the *M. robertsii* took place approximately 1.72 million years ago (copy a) and 0.214 million years ago (copy b). Thus, the divergence of the two copies in *M. guizhouense* from chrA in *M. robertsii* occurred much more recently than the speciation event between the two fungal species 15.1 million years ago. This suggests that the observed divergence is likely the result of a more recent horizontal transfer of chrA. The direction of this horizontal transfer remains unknown. Given our current data we propose two scenarios: i) chrA was transferred twice from *M. robertsii* to *M. guizhouense* (1.72 MYA and 0.214 MYA), or ii) chrA was initially present in *M. guizhouense* and duplicated 1.72 MYA, with one copy subsequently transferred to *M. robertsii* 0.214 MYA. Of course, more complex scenarios that involve more steps or additional species (note that our phylogenetic analysis was restricted to 36 available sequences) might be possible but would require more assumptions. In conclusion, the presence of two copies of chrA in *M. guizhouense*, with different separation times from chrA in *M. robertsii*, implies either two horizontal chromosome transfers or a mixture of two biological mechanisms – thus, in either scenario, quite complex dynamics underlie this pattern.

### chrA and chrB differ in their composition from the rest of the genome

Accessory chromosomes in fungi often differ in their sequence characteristics compared to the core chromosomes, with lower gene density and higher density of transposable elements (1). These differences may indicate other evolutionary constraints for accessory chromosomes than for the core genome, such as a smaller effective population size (1). We observed a comparable pattern for the accessory chromosomes chrA and chrB in *M. robertsii*. chrA and chrB were enriched in their proportion of transposable elements (TEs), which constituted approximately 31-32% of all sequences in the accessory chromosomes, with 9.1% in the rest of the genome (Fig. 5 A). The proportion of genes on the accessory chromosomes was lower, with 22% compared to 29% in the rest of the genome. Taken together, TE and gene density resulted in a clear compositional separation of the two accessory chrA and chrB from the core chromosomes (Fig. 5 B). In addition, the TE composition (Fig. S6 A, B), and the codon usage of the genes (Fig. S6 C) located on the accessory chrA and chrB differed from that of rest of the genome. The gene-wise relative synonymous codon usage (gRSCU), which measures the bias in the use of synonymous codons, was significantly lower for genes located on chrA and chrB compared to the rest of the genome of *M. robertsii* R3-I4 as well as compared to those genes located on the rest of the genome of the R1-A ancestral strain (all pairwise Wilcoxon rank-sum test with BH correction, all p < 2 × 10^−16^). Taken together, these differences in TE composition and codon usage suggest that the accessory chromosomes were horizontally acquired by *M. robertsii*. Furthermore, the codon usage of chrA also differed from that of all genes in the *M. guizhouense* genome, which suggests that chrA was also horizontally acquired by *M. guizhouense*. Therefore, chrA may have been horizontally acquired by both *M. guizhouense* and *M. robertsii* from a third, currently unknown species.

**Fig. 5:**
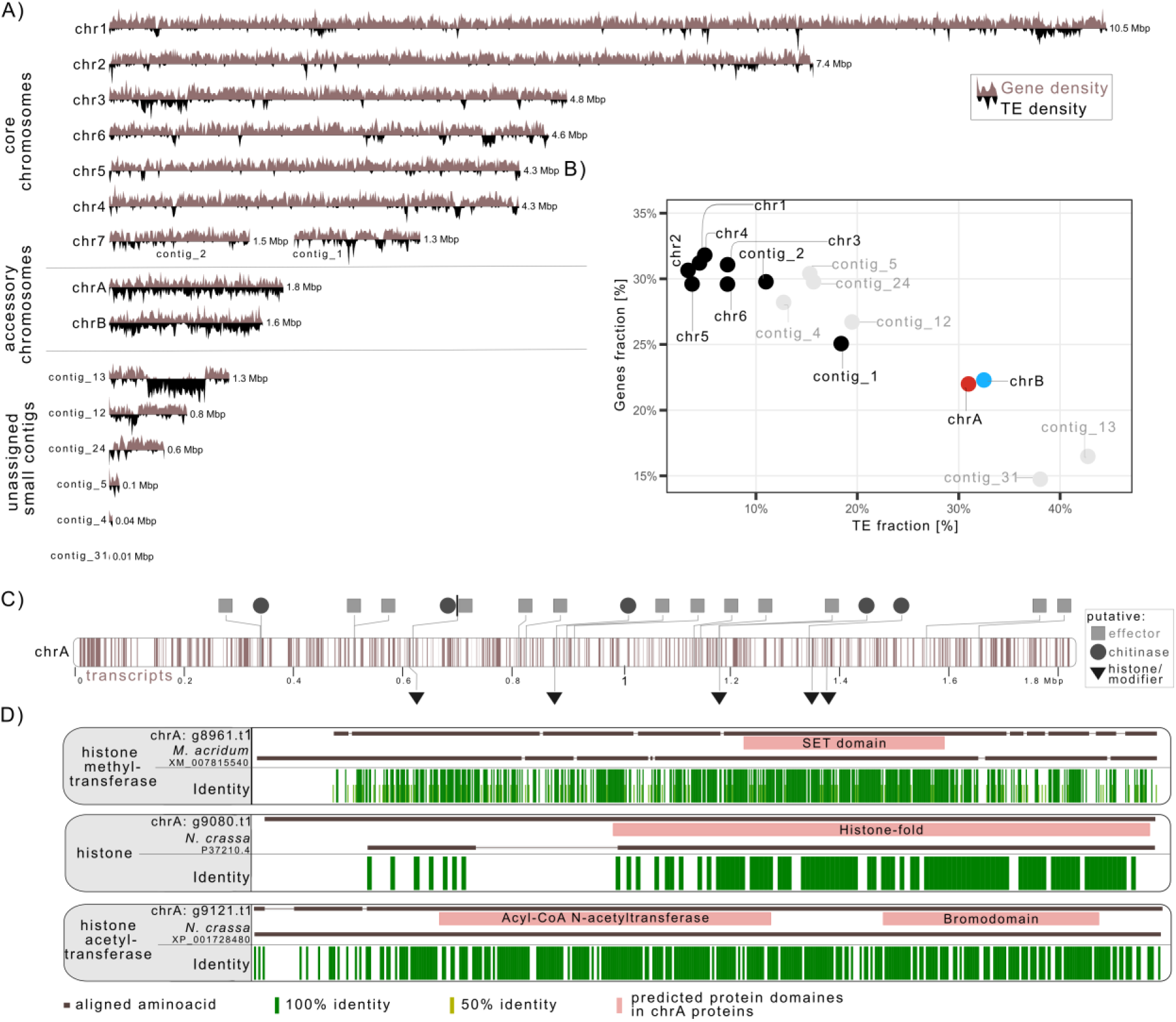
Overview of gene and transposable element annotation of the evolved *M. robertsii* R3-I4 strain. A) Karyoplot illustrating the distribution of genes (light brown) and transposable elements (TEs) (black) along the nanopore-based assembly of the evolved R3-I4 strain. B) Distribution of gene density vs TE density, highlighting the higher TE density and lower gene density observed in chrA (red) and chrB (blue) compared to other chromosomes (black) and unassigned small contigs (grey). C) Gene transcript distribution along chrA, with putative candidates that might influence either the interaction with the host or the chromatin conformation being indicated. D) Example alignments of proteins for one putative histone, and one putative histone methyl-transferase and one putative histone acetyltransferase with proteins of known function. Identical amino-acids are depicted in green.

### chrA encodes putative virulence factors and may influence its horizontal transfer

We were further interested if the proteins encoded by the genes on the accessory chromosomes could be potentially involved in the interaction with the insect host or the horizontal transfer of the chromosome and therefore functionally annotated the predicted transcripts. On accessory chrA, we could identify a total of 364 genes through *ab initio* annotation (Table S6, Table S7 A-B). Among them, 13 genes were predicted to encode putative effectors (i.e. small, secreted proteins that could potentially interfere with the host immune system) and ten genes were identified as Carbohydrate-Active Enzymes (CAZymes). Notably, three of these CAZymes were putative chitinases, suggesting a possible role in host insect cuticle degradation, which is a crucial step during the infection process, since the fungal spores need to penetrate the host’s chitin cuticle to be able to infect and eventually kill the host (31). Hence, the accessory chromosomes might be involved in the interaction with the insect host and by receiving the accessory chrA the recipient strain may have gained a fitness benefit by increased virulence.

On both accessory chrA and chrB, we further found genes that might be involved in the horizontal transfer of the accessory chromosomes. ChrA contained two genes encoding putative histone H2B proteins (g9016.t1, g9080.t1) and three genes encoding putative enzymes involved in histone post-translational modifications (g8961.t1, g9112.t1, g9121.t1) (Fig. 5 C). The two putative histone H2B proteins displayed significant identity in the histone-fold region with the histone H2B protein from the mold fungus *Neurospora crassa*. Among the three putative histone-modifying enzymes, two contained a SET domain specific to enzymes that methylate histone lysine residues, while the remaining gene showed high similarity to the histone acetyltransferase protein GCN5 (Fig. 5 D). Similarly, chrB harboured a gene encoding a putative histone H3 protein, which displayed high identity to the histone fold domain of *N. crassa* histone H3 protein (g10192.t1, aa44-136, 93% identity, accession XP_956003.1). ChrB also contained two genes encoding putative histone-modifying enzymes: a GCN5 acetyltransferase homolog (g10208.t1) and a SET domain-containing putative histone methyltransferase (g10199.t1). In all our observed intraspecies horizontal transfers of accessory chrA solely this chromosome and no other material was either transferred or retained. Hence there must be a distinguishing characteristic that separates chrA from the rest of the donor genome and also from the genome of the recipient strain. Histone modifications are known to be involved in many programmed DNA elimination in plants and animals (39). The presence of these putative histone proteins and histone-modifying enzymes on both chrA and chrB suggests their potential involvement in affecting the chromatin conformation and thereby potentially mechanistically influencing the horizontal transfer of the accessory chromosomes.

## Discussion

In this study, we describe the horizontal transfer of an entire accessory chromosome (chrA) between asexual strains and species of common insect-pathogenic fungi. We show that: i) two accessory chromosomes are present in *M. robertsii* and that one of them is frequently transferred horizontally during experimental co-infection between different *M. robertsii* strains. This transfer event involved only the accessory chromosome, while no other genetic material was transferred. ii) Although horizontal transfers occurred frequently, the transferred accessory chromosome was only able to outcompete the same strain lacking it, and to spread in the pathogen population under one experimental condition – infection of single ant hosts, but not when infecting ants accompanied by nestmates that provided social immunity. Therefore, the accessory chromosome alters the competitive ability of the recipient strain, and whether or not this results in fitness benefits is condition-dependent. In social ants, the fitness benefits provided by chrA when infecting single hosts were negated when colony members provided additional social immunity. iii) Importantly, we show that this horizontal transfer is not restricted to exchange within the same species in experimental co-infections, but also occurs naturally across species under field conditions, as the same accessory chromosome has been transferred between different species of *Metarhizium* much more recently than their speciation time of 15.1 MYA. Such horizontal transfer between different species could therefore be a possible mechanism for the spread of accessory chromosomes to new species.

Our characterization of multiple field-isolated strains of *M. robertsii* showed that this entomopathogen contains two accessory chromosomes with presence/absence polymorphism. To our knowledge, this is the first description of accessory chromosomes in pathogenic fungi of animals, where their presence and role is so far underexplored (40), in contrast to their common occurrence in plant-pathogenic fungi, where they have demonstrated effects on fitness and/or host range (1, 3). Notably, *M. robertsii*, like many other *Metarhizium* species, is – in addition to its parasitic lifestyle in insects – also closely associated with plants, in particular their roots. Therefore, the accessory chromosomes in *M. robertsii* may also be involved in the interaction with plants. Similar to accessory chromosomes in other fungal pathogens the accessory chromosomes in *M. robertsii* (chrA and chrB) differed in TE and gene content as well as codon usage from the rest of the genome, which could indicate different evolutionary histories or constraints in accessory vs core chromosomes.

In our experiment the chrA was transferred at least five times independently from *M. robertsii* R1 to the *M. robertsii* R3 strain, while no other genetic material was exchanged. While horizontal transfer of accessory chromosomes has been reported *in vitro* for the plant pathogens *Colletotrichum gloeosporioides* (10, 12), members of the *Fusarium oxysporum* species complex (7, 8, 13, 41) and possibly in *Alternaria alternata* pathotypes (42), the low frequency of these transfers necessitated the use of selectable fungicide resistance markers in these studies to identify and isolate those fungal cells, in which a horizontal transfer had occurred. Consequently, to date no horizontal transfer of chromosomes has been observed for these fungal pathogens during infection of their respective host (10, 12, 13, 41). In contrast, our study observed frequent horizontal transfer between *M. robertsii* strains during co-infection of ants, which represent a natural host of multiple *Metarhizium* species (31). This disparity suggests that the higher frequency of horizontal transfer observed in our study may in principle be attributable to parasexual-like processes described for species of the genus *Metarhizium* (43, 44), which is thought to lead to exchange of genetic material between co-infecting strains when both grow within the same host individual. However, the parasexual-like cycle in *Metarhizium* is thought to involve cell fusion resulting in cells with two different nuclei (heterokaryons), followed by karyogamy to produce unstable diploids that randomly lose chromosomes to regain haploidy (44). In our study, in contrast, we observed specific transfer of exactly one chromosome (chrA) without random genetic exchanges or loss of additional chromosomes. Thus, we propose that the observed horizontal transfer of chrA cannot be explained by the established parasexual-like processes alone. Instead, alternative mechanisms for accessory chromosome transfer must be in place. These could include either the transfer of accessory chromosomes between nuclei during the heterokaryon stage (12) or the selective degradation of all chromosomes except the one chromosome that was successfully transferred (12, 13). A process similar to the former has been described in *Saccharomyces cerevisiae*, where a mutant defective in nuclear fusion (kar1-1) generates transient heterokaryons, allowing chromosome transfer between nuclei before one of the nuclei is lost. This process, known as chromoduction, enables the transfer of whole chromosomes into a new genomic background (24, 45–47). A process similar to the latter (i.e. degradation of all but the transferred chromosomes) has been described as programmed DNA elimination in several plants and animals (reviewed in (39)), with the retained DNA sequences differing from the eliminated ones through DNA methylation, histones, or post-translational histone modifications, particularly in centromeric regions (39). We speculate that such a mechanism may also act during the horizontal transfer of chrA and possibly also chrB in *M. robertsii*. Since both accessory chromosomes contain genes encoding putative histones and histone-modifying enzymes, these may be involved in ensuring the transfer of only the accessory chromosomes. We hypothesize that the distinct chromatin conformations of the accessory chromosomes compared to the rest of the donor genome, allows either their preferential transfer to the recipient nucleus (similar to chromoduction), or evasion of degradation (similar to the PSR chromosome in the parasitoid wasp, *Nasonia vitripennis*, where the presence of the PSR chromosome affects the post-translational methylation of at least three histones in the genome that later becomes eliminated (48, 49)).

In addition to the horizontal transfer of chrA between strains of *M. robertsii* we show phylogenetic support of a past horizontal chromosome transfer between two different species of the genus *Metarhizium* in the field. While interspecific chromosome transfer has been proposed as a potential mechanism for how species can gain new accessory chromosomes, no such transfer had been previously documented to our knowledge. Existing experimental evidence on horizontal transfers has focused on different pathovars (e.g., *A. alternate*), biotypes (e.g., *C. gloeosporioides*), or formae speciales within species complexes (e.g., *F. oxysporum*) (6–8, 10, 13, 41, 42). Although these studies provide insights into the distribution of accessory chromosomes within species, they fail to account for their spread between species. The scarcity of fungal accessory chromosomes that exhibit synteny with each other in different species was considered as evidence against the hypothesis that these chromosomes had been acquired through horizontal transfer (2) and evidence for their origin by endogenous processes (i.e. from within the genome, e.g. via degenerative breakage, duplication, missegregation or Robertsonian chromosome fusion (2, 50, 51)). Here, we report exactly such a syntenic accessory chromosome being present in two species (*M. robertsii* and *M. guizhouense*). Moreover, the codon usage pattern of the accessory chrA differed from the rest of the genome of both *M. robertsii* and *M. guizhouense*. This suggests that it might even have originated from a third species and was then horizontally acquired. Taken together, we here show intra- and interspecific horizontal transfer of an accessory chromosome, supporting that horizontal chromosome transfers present a more common mechanism for a species to acquire novel accessory chromosomes than previously thought.

Our experimental study further gives unprecedented details into the dynamics and constraints of the spread of an accessory chromosome after its horizontal transfer into a new pathogenic strain. In our experiment chrA was horizontally transferred in five of the total 20 independent replicates. Strikingly, all three cases, where the strain that horizontally-acquired chrA outcompeted the strain lacking chrA were observed during infection of single ants. The two instances where the strain that horizontally-acquired chrA was outcompeted by the strain that had not acquired it, leading to its extinction, was observed during infection of ants in the presence of nestmates that provide social immunity. Hence, even when transfer occurs frequently, whether or not the strain harboring the acquired accessory chromosome will be able to establish itself in the pathogen population is a condition-dependent evolutionary process. However, a positive fitness effect may also not be required in all cases, as the accessory chromosome may still spread even in the absence of a benefit, purely due to a preferential transfer mechanism similar to a meiotic chromosome drive (52). It is tempting to speculate that since chrA and chrB contain genes that could influence their preferential transfer - like the putative histone and histone-modifying enzymes - that these accessory chromosomes could also be selfish genetic elements, manipulating and propagating their horizontal transmission.

In conclusion, we here describe accessory chromosomes with the potential to spread both within and across species of fungal pathogen species of the genus *Metarhizium*, which comprises many important insect-pathogenic species. By describing the characteristics of this novel accessory chromosome and its transfer and spread dynamics, we find horizontal transfer to be a very important mechanism in shaping the evolution of accessory chromosomes and whole genomes. Based on its preferential horizontal transfer chrA might be able to spread through the recipient pathogen populations in a process highly reminiscent of the spread of selfish genetic elements.

## METHODS

### Fungal strains

We used the six ancestral strains and the evolved lines of the fungal pathogens *Metarhizium robertsii* and *M. brunneum* obtained during the selection experiment performed by Stock et al. (32). In this experiment, the six ancestral strains were mixed in equal amounts and used to experimentally co-infect Argentine ants, which subsequently were either kept alone (individual treatment (abbreviation: I)) or in the presence of two nestmates (social treatment (abbreviation: S)) over ten consecutive host infection cycles (passages). An outline of the experiment is shown in Fig 1A, and details described in Stock et al. (32) and Supplementary Text S1. The six ancestral strains included three *M. robertsii* strains (R1-A: KVL 12-36 (C17), R2-A: KVL 12-35 (E81) and R3-A: KVL 12-38 (F19)) and three *M. brunneum* strains (B1-A: KVL 13-13 (G39), B2-A: KVL 12-37 (J65), and B3-A: KVL 13-14 (L105); all obtained from the University of Copenhagen, Denmark), which had all been isolated from the same field population and characterized by Steinwender et al. (53). For the evolved lines (32), we focused particularly on *M. robertsii* R3, for which we determined overall strain proportion and integration of chrA over the course of the experiment (at passages P1, P3, P5 and P10). *M. guizhouense* ARSEF977 was obtained from the ARS Collection of Entomopathogenic Fungal Cultures, Ithaca NY, USA. An overview of previously published reads and assemblies that were included in this study is given in Table S4.

### Molecular analysis of fungal strains

We performed both long read sequencing (Nanopore and PacBio) as well as short read sequencing (Illumina) for ancestral and evolved strains, as well as microsatellite analysis for spore strain identification and chrA presence/absence determination, as detailed in Supplementary Text S1. Nanopore sequencing was performed by the Next Generation Sequencing Facility at Vienna BioCenter Core Facilities (VBCF), member of the Vienna BioCenter (VBC). PacBio sequencing of *M. guizhouense* was performed at the Max Planck Genome Centre Cologne, Germany using Sequel IIe (Pacific Biosciences). Illumina Sequencing was performed at Eurofins Genomics GmbH (Ebersberg). An overview of the sequencing reads generated in this study is given in Table S8.

### Bioinformatic analysis

The Bioinformatic steps are explained in the Supplementary Text S1.

### Statistical data analysis

All statistical analyses were conducted in R (version R3.6.0) using the suite R Studio (1.2.1335), as described in more detail in Supplementary Text S1. A summary of all statistical test results is given in Supplementary Data S1.

### Data availability

Sequencing reads have been deposited in the Sequence Read Archive and are available under the BioProject PRJNA1017668. The Nanopore-based assemblies and Gene and TE Annotations were deposited at NCBI under the BioProjects PRJNA1015426, PRJNA1015429, PRJNA1015431.

All source data for the figures are given in the source data file (Supplementary Data S1). Note that of the total of 2626 spore-clones shown in Fig. 2, 872 had been produced, analyzed for strain identity and previously published in Stock et al. (32). The production and strain-identification of the remaining 1754 spore-derived clones, as well as the presence/absence characterization of chrA within all 883 R3 spore-clones were performed in this study.

## Acknowledgements

We thank Bernhardt Steinwender, Jorgen Eilenberg and Nicolai V. Meyling for the fungal strains. We further thank Chengshu Wang for providing the short sequencing reads for *M. guizhouense* ARESF977 he used for his published genome assembly, and Kristian Ullrich for help in the bioinformatics analysis for methylation pattern in Nanopore reads, and the Vienna BioCenter and the Max Planck Society for the use of their sequencing centers. We thank Barbara Milutinović and Hinrich Schulenburg for discussion, and Tal Dagan and Jens Rolff for comments on a previous version of the manuscript. Fig1 A was created with BioRender.com.

This study received funding by the European Research Council (ERC) under the European Union’s Horizon 2020 Research and Innovation Programme (No. 771402; EPIDEMICSonCHIP) to S.C. and by the German Research Foundation (DFG grant HA9263/1-1) to M.H.

## Supplementary figures

**Fig S1.**
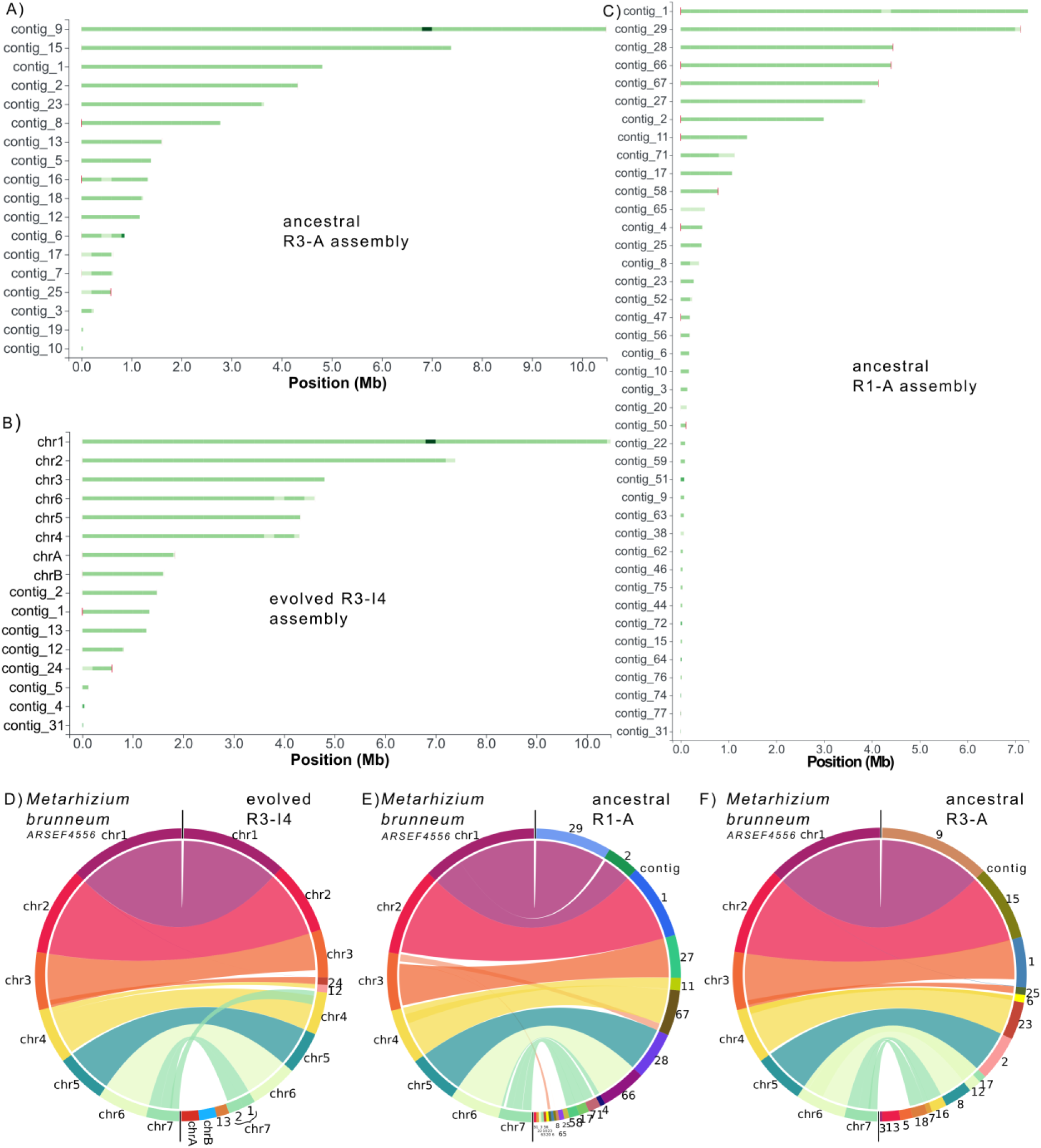
Nanopore-based assemblies of *Metarhizium robertsii* ancestral R1-A, ancestral R3-A and the adapted R3-I4 strains at near chromosome level. Tapestry reports of the nanopore-based assemblies of A) ancestral R3-A, B) adapted R3-I4 and C) ancestral R1-A strain. Red marks represent the presence of telomere repeats, where the intensity of the red colour is proportional to the number of repeats detected. Green intensity is proportional to coverage. For a detailed description, see also Table S1. D) Synteny between the nanopore-based assemblies of *M. robertsii* strains R3-I4, R1-A and R3-A generated in this study and the *M. brunneum* ARSEF4556 reference assembly (GCA_013426205.1) (33). Note that for R3-I4 chromosomes were labelled based on their synteny to the *M. brunneum* reference assembly.

**Fig. S2.**
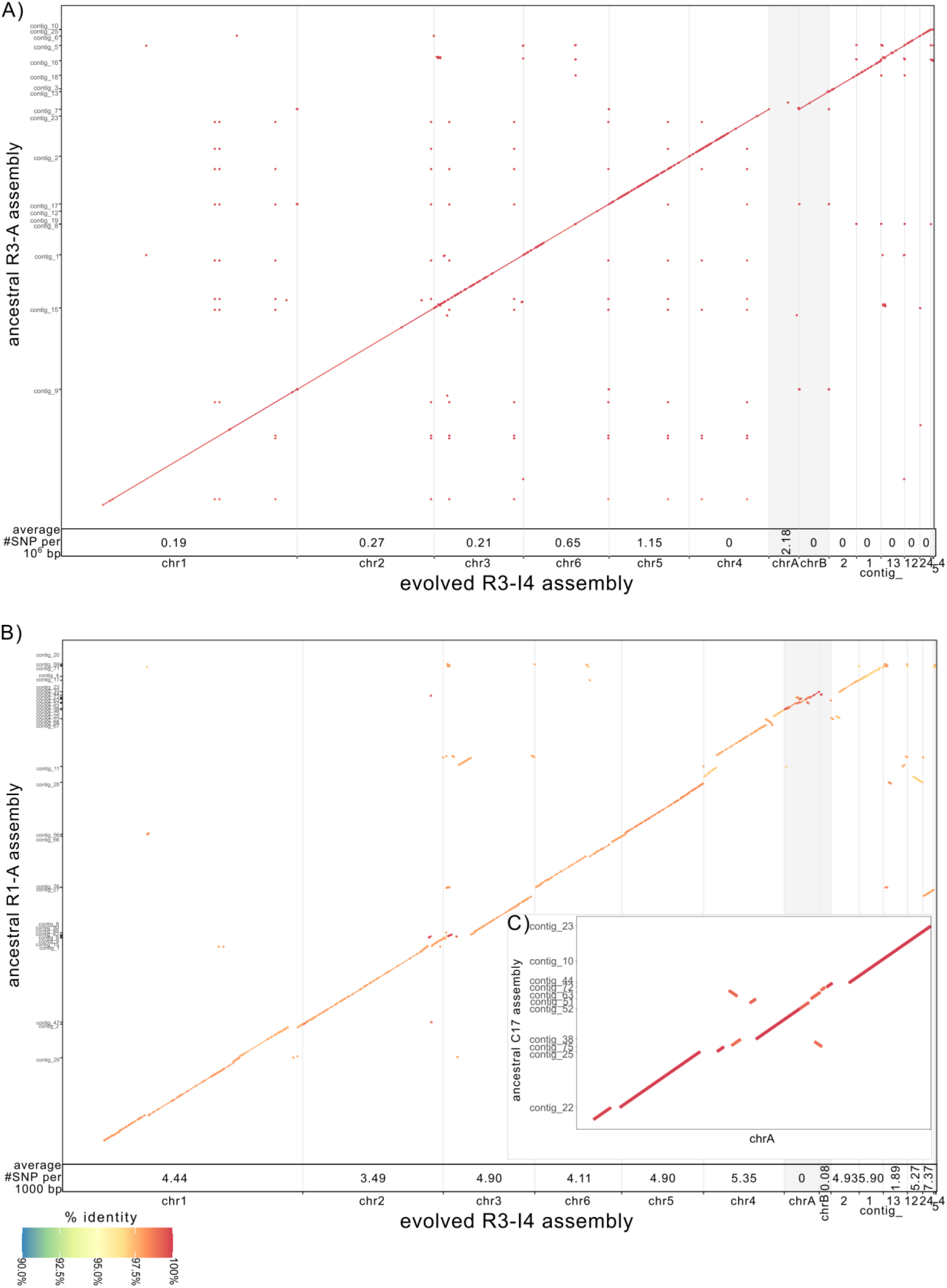
Synteny plot between the nanopore-based assemblies of the A) ancestral R1-A or B) ancestral R3-A with the evolved R3-I4 strain, with synteny for chrA and chrB of the R3-I4 highlighted in grey shade. In C) the alignment of R1-A contigs syntenic with chrA of the evolved R3-I4 is shown. The SNP density based on these alignments for each of the contigs is given at the bottom of each graph.

**Fig S3:**
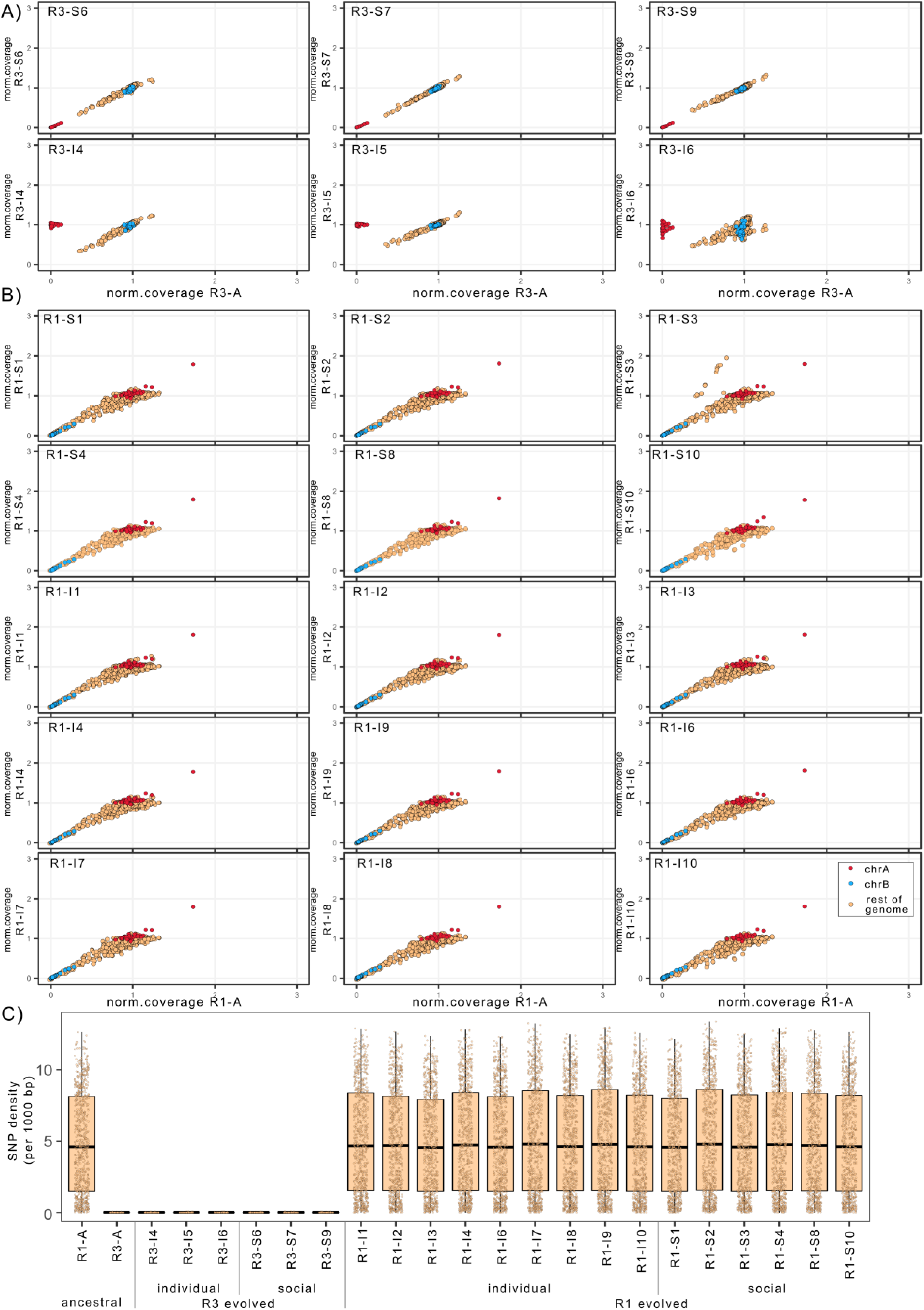
Coverage analysis and SNP/InDel distribution failed to detect horizontal transfer of genetic material in addition to chrA. Illumina sequence coverage analysis of the A) adapted R3 strains compared to the coverage of the ancestral R3-A strain or the B) adapted R1 strains compared to the coverage of the ancestral R1-A strain. With the exception of chrA for the individual-adapted R3 strains, no change in sequence coverage was detected. C) Distribution of SNPs/InDels that are specific to the R1-A (present in the R1-A but absent in the R3-A). No 50 kb windows in the adapted R3 strains showed increased SNP density. Hence no large-scale transfer of genetic material, in addition to chrA, occurred from the ancestral R1-A strain to the adapted R3 strains. In the adapted R1 strains, there was no change in the distribution of SNPs/InDels compared to the ancestral R1-A strain, indicating that no large-scale transfer of genetic material to the adapted R1 strains occurred. Note that the rDNA cluster was excluded from the analysis for visual clarity due to its high coverage.

**Fig. S4:**
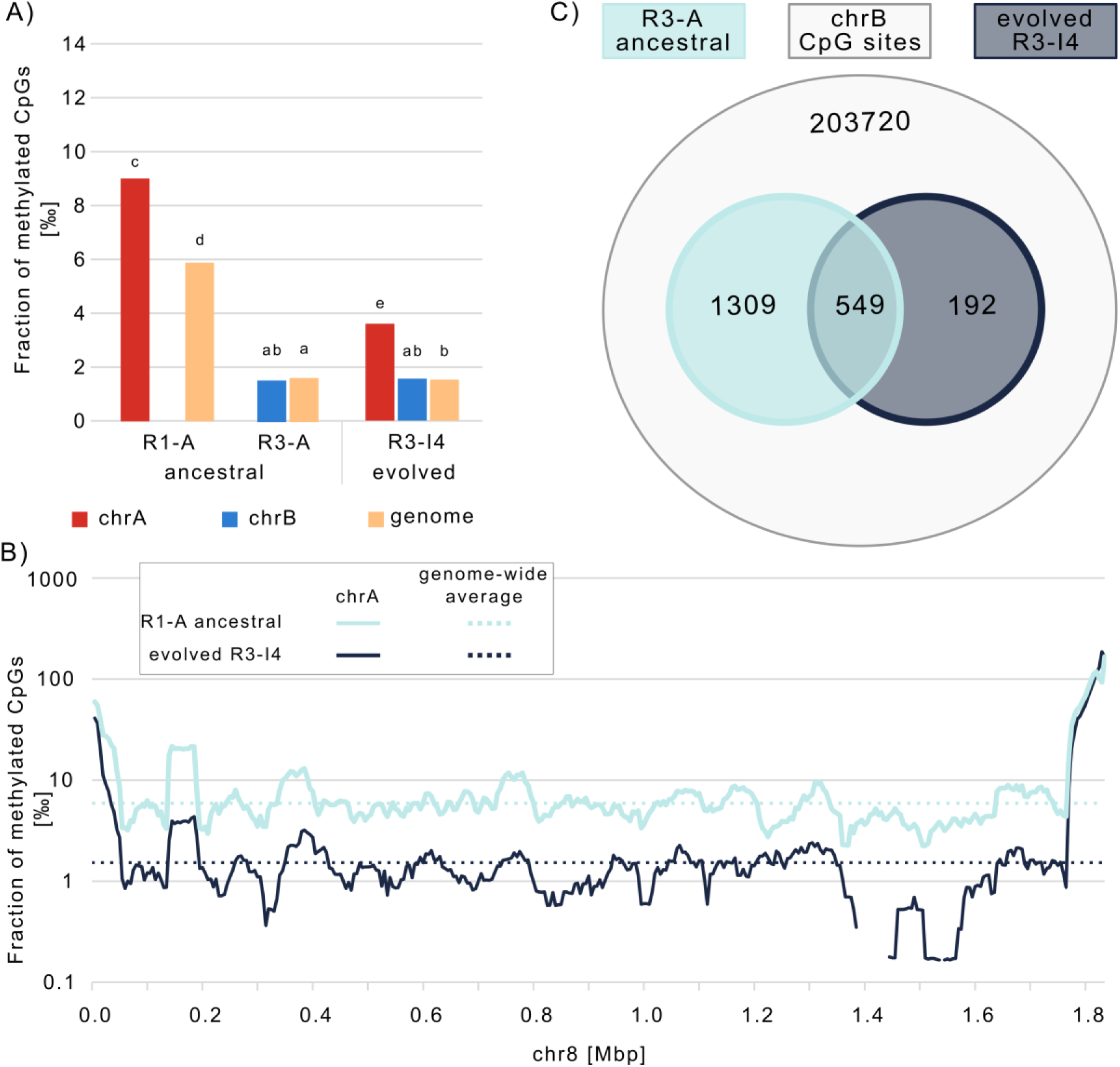
The methylation pattern differed between ancestral R1-A and R3-A strains. ChrA retained part of the ancestral methylation pattern after horizontal transfer from R1-A to R3-A. A) Fraction of methylated cytosines in CpG contexts for chrA, chrB and the rest of the genome in the ancestral R1-A and R3-A strains and the adapted R3-I4 strain. ChrA showed higher methylation in both the ancestral R1-A strain and the adapted R3-I4 strain than chrB and the rest of the genome (identical letters above groups indicate non-significance at α<0.05 determined by Fisher exact test with BH-adjustment for multiple testing). B) Fraction of methylated cytosines in CpG contexts in 50kb windows (sliding: 5 kb) along chrA for the ancestral R1-A (turquoise) and evolved R3-I4 (dark-grey) strains. C) Venn diagram showing the overlap of highly methylated CpG sites (>25% methylation) of chrA (total number CpG sites 203720) between the ancestral R1-A and the evolved R3-I4 strain.

**Fig. S5.**
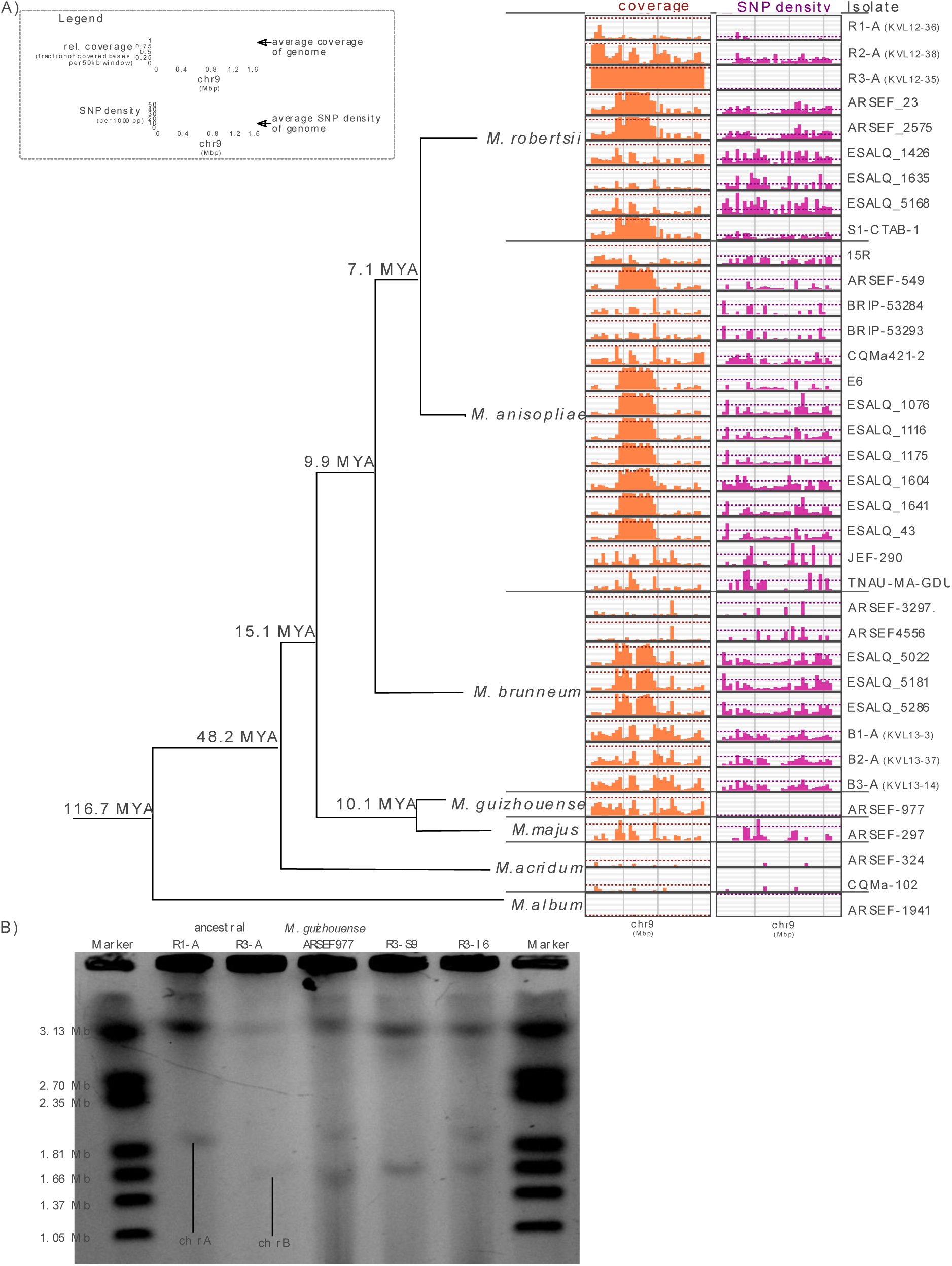
Presence/absence polymorphism of chrB in published genomes of species of the genus *Metarhizium* and pulsed-field gel electrophoresis showing the presence of small chromosomes similar in size to chrA and chrB in *M. guizhouense*. A) Phylogeny and distribution of relative sequence coverage (fraction of bases covered in 50 kb windows) in orange and SNP density per 1000 bp in 50 kb windows in pink, with respective genome-wide averages shown as dotted lines. Note: TEs were excluded from the analysis. Phylogeny adapted from published Hu and colleagues, 2014 (38). MYA: Million years ago. B) PFGE of *M. guizhouense* in comparison to the ancestral *M. robertsii* R1-A and R3-A, as well as two evolved R3 strains R3-S9 (lacking chrA) and R3-I4 (including chrA).

**Fig. S6:**
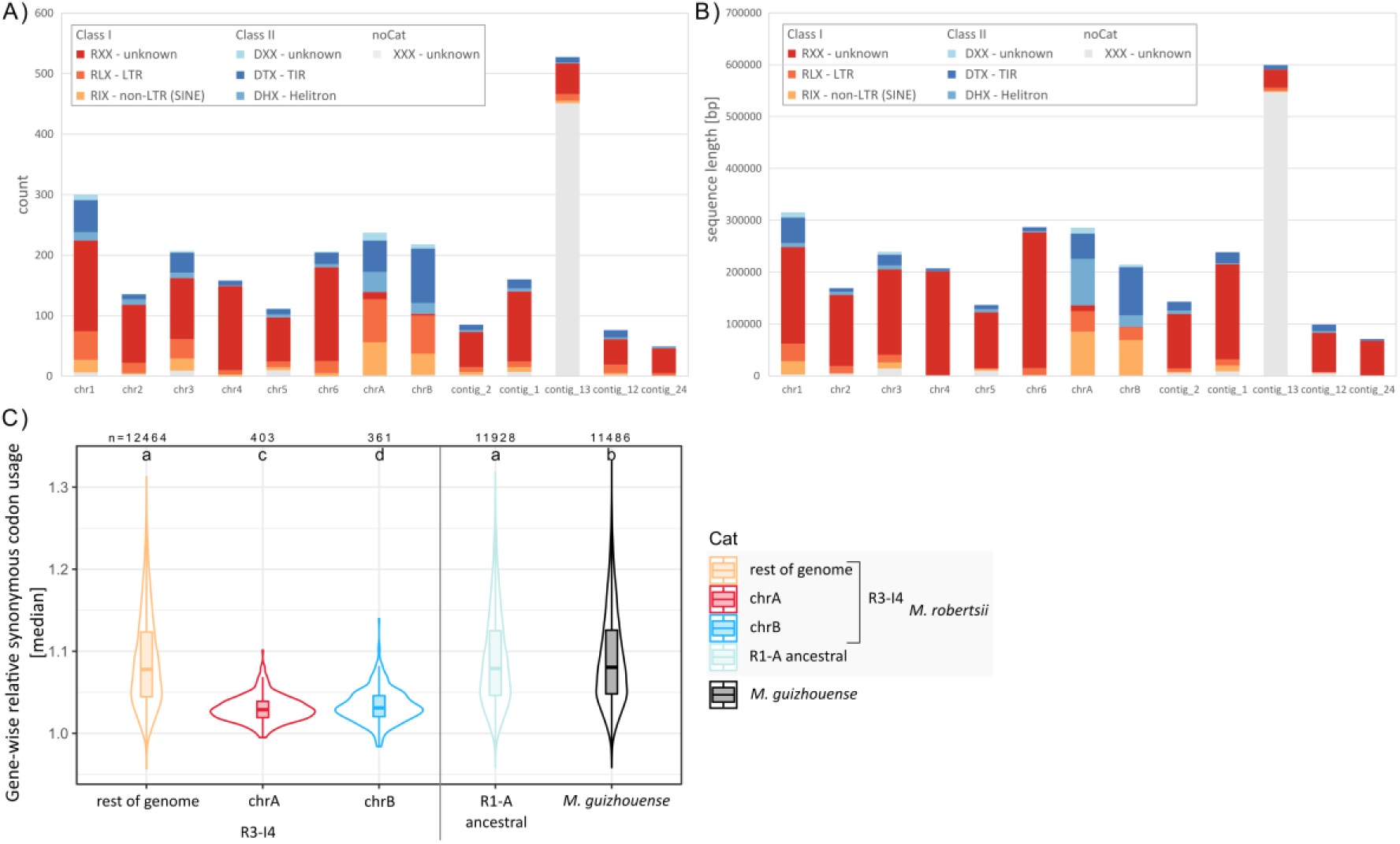
TE composition and codon usage of the accessory chromosomes chrA and chrB differed from the rest of the genome. A) Number and B) cumulative sequence length of the indicated TE classes and orders of the *M. robertsii* R3-I4 strain. ChrA and chrB have a higher proportion of Class I retrotransposons of LTR and SINE order and virtually lack the unknown order of the Class I retrotransposons that dominates (dark-red) in the other chromosomes. C) Gene-wise relative synonymous codon usage for genes located on chrA and chrB compared to genes located on the rest of the R3-I4 genome and R1-A and the *M. guizhouense* ARSEF-977 genomes. Identical letters above the individual plots indicate that the respective groups were not significantly different (pairwise Wilcoxon rank-sum tests with BH correction, α=0.05).

## Supplementary Tables

Table S1: Comparison statistics of Nanopore-based assemblies

Table S2: Number of SNPs and small InDels compared to R3-I4 that were not already present in the ancestral R3-A strain

Table S3: Number of methylated Cytosines in CpG context

Table S4: Overview of previously published Assemblies and Reads included in this study Table S5: Number and phases of SNPs and InDels in *M. guizhouense* on the R3-I4 assembly. Table S6: Comparison of genome annotations

Table S7 A: R3-I4: Annotation of chrA

Table S7 B: R3-I4: Genome-wide annotation

Table S8: Overview of sequencing information generated within this study

## Supplementary Data

Data S1: All source data of figures and exact p values for statistical tests

## Supplementary Text

Text S1: Detailed Methods and Materials

