## Supplemental Text S1 for "Frequent horizontal chromosome transfer between asexual fungal insect pathogens"

**Text S1.** Detailed methods and materials

### 1) Fungal strains and selection experiment outline

This study analyzed 30 fungal strains of the entomopathogenic fungal genera *Metarhizium robertsii* and *brunneum,* six of which had been used as the starting strains of a selection experiment by Stock et al. (1) in ant hosts (ancestral strains), and 24 of which represent the evolved strains at the end of the experiment. The six starting strains (3 *M. robertsii* strains KVL 12-36 (C17), KVL 12-38 (F19), and KVL 12-35 (E81) and 3 *M. brunneum* strains KVL 13-13 (G39), KVL 12-37 (J65), and KVL 13-14 (L105; all obtained from the University of Copenhagen, Denmark (B. Steinwender, J. Eilenberg and N.V. Meyling)) had been collected from an agricultural field in Denmark by Steinwender et al. (2). After being grown as monospore cultivar, the six strains were mixed in equal amounts (total concentration of 1 × $10^6$ spores $ml^{-1}$) and were used to co-infect workers of the Argentine ant, *Linepithema humile*, over ten host infection cycles in each of two selection treatments (each in ten independent replicate lines), as detailed in Stock et al. (1). In the “individual treatment”, the worker ant remained alone after exposure, whilst it was accompanied by two untreated nestmates in the “social treatment”. After each infection cycle, the spores growing out of the first eight dying workers per replicate line (ants dying within the first 24 h after exposure, as well as nestmates were not considered) were harvested, mixed and used to infect a new round of hosts, which were then consistently kept either under the individual resp. social treatment conditions depending on the replicate. After passage 5 and 10 of the experiment, it was determined, which strains were still present in the mix (Stock et al., Fig. 1). At the end of the experiment, 16 lines contained only a single strain genotype (identified by microsatellite analyses) whilst four lines were a mix of two persisting spore types from two different starting strains (see Stock et al. (1)). One spore of each type of these evolved lines was expanded to obtain monospore cultivars of all the strains that had been able to persist after the ten host passages. Phenotypic analyses of the lines (16 single-strain lines and the four 2-strain lines, mixed in the proportion of strain presence at the end of the experiment) found that the lines that had adapted to only the individual immune defenses of their single ant hosts showed increased virulence, whilst the lines adapted to the social immunity of the ants kept in groups, showed increased production of spores with a reduced content of the fungal cell membrane compound ergosterol (1).

In this study, we Illumina-sequenced these 24 evolved strains, as well as their six ancestral strains. We follow the terminology used in Stock et al. (1), in that the three *M. robertsii* starting strains are named R1 to R3 (R1: KVL 12-36 (C17), R2: KVL 12-38 (F19), R3: KVL 12-35 (E81)) and the three *M. brunneum* strains B1-B3 (B1: KVL 13-13 (G39), B2: KVL 12-37 (J65), B3: KVL 13-14 (L105)). To distinguish them from the evolved strains at the end of the experiment, we here extend their numbering by an “-A” for “ancestral, i.e. R1-A, for example, representing the ancestral strain of R1. The evolved strains are numbered following their replicate line number (Fig. 1, with strain identity (given by microsatellite identification performed in Stock et al.), using “I” for the “individual” and “S” for the “social” treatment, i.e., as an example, the strain that was identified as R3 in the individual treatment replicate line 4, is abbreviated as R3-I4.

In addition to the analysis of the ancestral and evolved lines at the end of the experiment, we were here interested in the spore diversity within our evolved lines at different time points of the experiment, that is after passage 1, 3, 5 and 10. To this end, we plated the stored spore suspensions, picked single-clone colonies and determined their strain identity and the presence / absence of chrA, as detailed below. This was done for all evolved lines that showed R3 being present in passage 5, independent of whether it succeeded into passage

10 or not. Therefore, we analyzed replicates R3-I4, R3-I5 and R3-I6 from the individual treatment and R3-S1, R3-S6, R3-S7, R3-S8, R3-S9 for the social treatment.

### 2) Pulsed-field gel electrophoresis (PFGE)

Spores were grown in liquid LB medium (2% sucrose, 1% peptone, 0.3% yeast extract and 0.5% NaCl [w/V]) at 23°C, 200 rpm for 5-10 days, filtered through a sieve, centrifuged (3000 x g, 10 min, RT), the pellet was washed and centrifuged again in 1 X TE buffer. The pellet was resuspended in 500 µl $H_2O$ and 500 µl 2.2% low melting agarose and filled into plug casts. After solidification, ten plugs were incubated in 5 mL lysis buffer (0.45 M EDTA, pH 8.0, 1% SDS, 1.5 mg/mL Proteinase K) for 24 h at 55°C. After 24 h, the lysis buffer was replaced with fresh lysis buffer and the incubation was repeated. After 48 h, the plugs were washed three times with 1x TE for 20 min each and stored in 0.5 M EDTA. PFGE was performed in 1x TBE buffer at 14°C, 3 V/cm, 106° and 250-1000s switching time in a CHEF Dr III system using *Hansenula wingei* chromosomes (1.05-3.13 Mb) as size marker (Bio-Rad, Hercules, CA, USA).

### 3) Proportion of chrA-containing spores over the course of the experiment

The spore suspensions of passages 1, 3, 5 and 10 from the eight replicate lines that contained the R3 strain at least until passage 5 of the experiment (see Fig. 1 of Stock et al. (1)), were plated on selective medium agar plates, containing 6.5% Sabouraud dextrose agar (Sigma-Aldrich), 1 ml each of Syllit 450 SC (110 mg/ml; Kwizda), Chloramphenicol (100 mg/ml; Sigma-Aldrich) and Streptomycin (100 mg/ml; Sigma-Aldrich) and incubated for one week at 23°C in the dark. Individual clones were selected and transferred to DNeasy 96-well plates (Qiagen) containing 50 µl of nuclease-free water (Sigma-Aldrich). Samples were homogenized in a TissueLyser II (Qiagen) using approx. 100 mg of glass beads (425-600 µm; Sigma) in two steps (2 × 2 min at 30 Hz). Total DNA of the spore-cultivars was extracted using DNeasy 96 Blood and Tissue Kit (Qiagen) according to manufacturer's instructions, with a final elution volume of 50 µl Buffer AE. R3 clones were identified using the same microsatellite loci as in Stock et al. (1) Ma307 and Ma2054. Presence of chrA was determined in all R3 spore clones using primers designed in this study. For primer sequences see Table S9. All reactions were performed using MyTaq HS Red Mix (Bioline), 10 pmol of each primer (Ma307 only 5 pmol per primer) and 4 µl of genomic DNA. PCR amplifications for strain identification were performed as follows: initial denaturation at 95 °C for 1 min, followed by 35 cycles of 30 s at 95 °C, 1 min at 59°C (Ma307) resp. 60 °C (Ma2054) and 1 min at 72°C and a final extension step at 72°C for 7 min. For the detection of chrA in R3 clones we used three sets of primers targeting different regions of the chrA sequence, to make sure that the whole chrA was transferred. In addition, the multiplex PCR contained a set of primers targeting a *Metarhizium robertsii* specific sequence to confirm sufficiently high DNA quality of the samples. Amplifications were performed as follows: initial denaturation at 95 °C for 1 min, followed by 30 cycles of 15 s at 95 °C, 15 s at 60 °C and 10 s at 72°C. In total 2626 individual spore clones (872 of which were already published in Stock et al. (1) and 1754 of which were produced in the course of the current study) were used. 883 of these clones were found to be R3 and were further analyzed for the presence of chrA. A detailed summary of the results can be found in Suppl. Data S1.

Table S9: Primers used within this study

| Aim | Primer name | Sequence (5' → 3') | Product size [bp] |
|---|---|---|---|
| R3 strain identification | Ma307_F<br>Ma307_R | CATGCTCCGCCTTATTCCTC<br>GGGTGGCGAAGAAGTAGACG | 161 |
| | Ma2054_F<br>Ma2054_R | GCCTGATCCAGACTCCCTCAGT<br>GCTTTCGTACCGAGGGCG | 230 |
| detection of chrA | ChrA_left_F<br>ChrA_left_R | ACTTGCCGCTCGAAAAACAC<br>CCTATTGGACCTTCCGTGCAA | 150 |
| | ChrA_center_F<br>ChrA_center_R | AAAGATCGCGGGTGACAACT<br>AACAGGACATCACGCTCTGC | 200 |
| | ChrA_right_F<br>ChrA_right_R | ACAGGCACGTGAGCTTCTAC<br>GTTCTAGCAGTGGTTGACGGA | 250 |
| | Mrobertsii_F<br>Mrobertsii_R | GACGAAAAGATTTGCGGCACC<br>TCACGATCCTTGAGACACGC | 116 |

**4) Statistical tests**

Statistical tests were performed on the methylation data (Fig. S4 A) using a two-sided Fisher exact test comparing the count data for the two indicated groups for the total number of CpG sites and the number of methylated sites. The p-values were adjusted for multiple testing using the BH correction. To compare codon usage bias (Fig. S6 C), all groups were first compared using a Kruskal-Wallis rank-sum test and later pairwise comparisons were calculated using the Wilcoxon rank-sum test (for unpaired data). The exact p-values can be found in Supplementary Data S1.

**5) Generating Genome Assemblies**

**DNA preparation for sequencing**

DNA for Nanopore and PacBio sequencing was extracted using the Blood & Cell Culture DNA Midi Kit (Qiagen, Hilden, Germany). Cells were grown in LB media at 23°C, 200 rpm for five to seven days and cells were collected by centrifugation (2000xg, 10min). For Nanopore und Illumina sequencing the fungal material was harvested by vacuum filtration (Filtermax filter top 0.22 µm, 500 mL). The cells were washed twice using ddH2O and afterwards freeze dried using a FreeZone 2.5 Liter Benchtop Freeze Dry System (Labcono) with the following settings: -50°C, 0.04 mbar and 23°C plate-temperature, ~ 18 hours. For PacBio sequencing the material was harvested by centrifugation (10 min, 3000 x g). Cells were ground to a fine powder in liquid nitrogen and 100 mg of the resulting powder was used for DNA isolation according to the manufacturer’s instructions. Nanopore sequencing was performed by the Next Generation Sequencing Facility at Vienna BioCenter Core Facilities (VBCF), member of the Vienna BioCenter (VBC), Austria. PacBio sequencing of *M. guizhouense* was performed at the Max Planck Genome Centre Cologne, Germany using Sequel IIe (Pacific Biosciences). Illumina Sequencing was performed at Eurofins Genomics GmbH (Ebersberg). An overview of the sequencing reads generated in this study is given in Table S8.

**Nanopore Sequencing**

**I**

**Sequencing: R9.4.1 Minion Flow cells. Base calling was performed using** guppy v5.0.11 using the model dna_r9.4.1_450bps_hac (high accuracy).

For the generation of the nanopore sequencing based assemblies we used the pipeline described in (5). In short:
Called Nanopore reads were filtered to reads longer than 5000 bases using NanoFilt (v2.3.0) (6).

```
gunzip -c reads.fastq.gz | NanoFilt  -l 5000 Reads.fastq.gz | gzip > Reads500.fastq.gz
```

Trimming of Illumina PE reads (trimmed using Trimmomatic V0.39  (7):

```
Trimmomatic  java -jar /trimmomatic-0.39.jar PE R1.fastq R2.fastq R1_paired.fastq R1_unpaired.fastq R2_paired.fastq R2_unpaired.fastq ILLUMINACLIP:TruSeq2-PE.fa:2:30:10:2:keepBothReads LEADING:3 TRAILING:3 MINLEN:36

cat R1_paired.fq.gz R1_unpaired.fq.gz R2_paired.fq.gz R2_unpaired.fq.gz > trimmed.fq.gz
```

Correction of Nanopore reads by trimmed Illumina reads using FMLRC 2 (0.1.4) (8).

```
gunzip -c trimmed.fq.gz | awk 'NR % 4 == 2' | sort | tr NT TN | ropebwt2 -LR | tr NT TN | fmlrc-convert comp_msbwt.npy

fmlrc2 comp_msbwt.npy Reads5000.fastq.gz Reads5000.corrected.fastq.gz
```

Further trimming of corrected reads using Canu (v2.1.1) (9).

```
canu -trim -p genomeID -d genomeID genomeSize=38m -corrected -nanopore Reads5000.corrected.fastq.gz
```

Assembly of corrected and trimmed Nanopore reads using flye (V 2.8.3) and two rounds of polishing (10)

```
flye --nano-corr Reads5000.corrected.trimmed.fastq.gz --out-dir outdir --threads 11 --trestle -i 2
```

Assembly polishing by uncorrected Nanopore reads using Racon (V1.4.20) (11) (two times):

Mapping of uncorrected Nanopore reads onto the assembly  using minimap2 version (2.22-r1101)

```
minimap2 -ax map-ont -t 6 -L canu5000_assembly.fasta 3000.Reads.fastq.gz > 5000Reads_on_canu5000.sam

racon -t 6 --no-trimming \

5000.Reads.fastq.gz \
```

```
5000Reads_on_canu5000.sam canu5000_assembly.fasta >
canu5000_assembly.racon1.fasta
```

Assembly polishing by uncorrected Nanopore reads using Medaka (V 1.4.3):

```
medaka_consensus -i 3000.Reads.fastq.gz \

-d canu5000_assembly.racon2.fasta \

-o outdir \

-t 4 -m  r941_min_high_g360
```

Assembly polishing by Illumina reads using Pilon (V 1.24) (12) after mapping with BWA-mem2 (V 2.2.1) (13) and samtools (1.12) (14) (4X):

```
bwa-mem2 -t 11 canu5000_assembly.racon2.medaka.fasta trimmed.fq.gz | samtools
sort Illumina_on_canu5000_assembly.racon2.medaka.pilon1.sorted.bam

java -Xmx16G -jar /pilon-1.24-0/pilon.jar --genome
canu5000_assembly.racon2.medaka.fasta \

--bam Illumina_on_canu5000_assembly.racon2.medaka.pilon1.sorted.bam \

--output canu5000_assembly.racon2.medaka.pilon\

 --outdir outdir --changes
```

The final assembly was analyzed using Tapestry (V 1.0.0) using the telomeric sequence TTAGGG (15)

```
weave -a canu5000_assembly.racon2.medaka.pilon4.fasta -r 5000Reads.fastq.gz \

 -t TTAGGG -o outdir -c 10
```

Due to the lower number of reads >5000 bases for the R1-A reads >3000bp were used for the Tapestry analysis

All contigs that showed a lower average coverage than 30 and larger than 100 were excluded from the assembly (this affected five contigs of the R3-I4  and 22 contigs of the R1-A and three contigs (130 kb) of the R3-A genome assembly .

The final statistics of the assembly are:

Table S10: Summary statistics of final genome assemblies (generated with Quast (V 5.0.2) (16).

| Statistics without reference | R1-A | R3-A | R3-I4 |
|---|---|---|---|
| # contigs | 41 | 18 | 16 |
| # contigs (>= 0 bp) | 41 | 18 | 16 |
| # contigs (>= 1000 bp) | 41 | 18 | 16 |
| # contigs (>= 5000 bp) | 40 | 18 | 16 |
| # contigs (>= 10000 bp) | 38 | 18 | 16 |
| # contigs (>= 25000 bp) | 36 | 16 | 15 |
| # contigs (>= 50000 bp) | 30 | 18 | 14 |
| Largest contig | 7267018 | 10479110 | 10465571 |

| | | | |
|---|---|---|---|
| Total length | 42768942 | 43163266 | 45009773 |
| Total length (>= 0 bp) | 42768942 | 43163266 | 45009773 |
| Total length (>= 1000 bp) | 42768942 | 43163266 | 45009773 |
| Total length (>= 5000 bp) | 42764321 | 43163266 | 45009773 |
| Total length (>= 10000 bp) | 42746527 | 43163266 | 45009773 |
| Total length (>= 25000 bp) | 42702419 | 43106281 | 44996207 |
| Total length (>= 50000 bp) | 42487955 | 43163266 | 44953661 |
| N50 | 4406720 | 4812484 | 4807498 |
| N75 | 2994606 | 2774337 | 4300840 |
| L50 | 4 | 3 | 3 |
| L75 | 7 | 6 | 6 |
| GC (%) | 49.76 | 50.29 | 50.24 |
| Mismatches | | | |
| # N's | 0 | 0 | 0 |
| # N's per 100 kbp | 0 | 0 | 0 |

### 6) Generation of Annotations

Transposable Element annotation was generated using the REPET pipeline (17, 18) following the pipeline as described in (19). In short, first a denovo annotation was created.

TEdenovo pipeline:

launch_TEdenovo.py -P R1-A -f MCL >& denovo.txt &

Output: R1-A_sim_denovoLibTEs_filtered.fa (filtered consensus sequences of TEs)

Second, this filtered consensus sequences of TEs identified by the denovo annotation were used to annote the TE in the genome. Here, two rounds of TEannot conducted.

First Round of TEannot using the sim_denovoLibTEs_filtered.fa as refTE.fa

TEannot.py -P R1-A_annot -C TEannot.cfg -S 1

TEannot.py -P R1-A_annot -C TEannot.cfg -S 2 -a BLR -v 2

TEannot.py -P R1-A_annot -C TEannot.cfg -S 2 -a RM -v 2

TEannot.py -P R1-A_annot -C TEannot.cfg -S 2 -a CEN -v 2

TEannot.py -P R1-A_annot -C TEannot.cfg -S 2 -a BLR -r -v 2

TEannot.py -P R1-A_annot -C TEannot.cfg -S 2 -a RM -r -v 2

TEannot.py -P R1-A_annot -C TEannot.cfg -S 2 -a CEN -r -v 2

TEannot.py -P R1-A_annot -C TEannot.cfg -S 3 -c BLR+RM+CEN -v 2

TEannot.py -P R1-A_annot -C TEannot.cfg -S 7 -v 2

Output: R1-A_annot_chr_allTEs_nr_join_path.annotStatsPerTE_FullLengthFrag.fa

Second Round of TEannot using the R1-A_annot_chr_allTEs_nr_join_path.annotStatsPerTE_FullLengthFrag.fa as FLF_refTE.fa

TEannot.py -P  R1-A_FLF  -C TEannot.cfg -S 2 -a BLR  -v 2

TEannot.py -P  R1-A_FLF  -C TEannot.cfg -S 2 -a RM -v 2

TEannot.py -P  R1-A_FLF  -C TEannot.cfg -S 2 -a CEN -v 2

TEannot.py -P  R1-A_FLF  -C TEannot.cfg -S 2 -a BLR -r -v 2

TEannot.py -P  R1-A_FLF -C TEannot.cfg -S 2 -a RM -r -v 2

TEannot.py -P  R1-A_FLF  -C TEannot.cfg -S 2 -a CEN -r -v 2

TEannot.py -P  R1-A_FLF -C TEannot.cfg -S 3 -c BLR+RM+CEN  -v 2

TEannot.py -P  R1-A_FLF -C TEannot.cfg -S 7 -v 2

PostAnalyzeTELib.py -a 3 -g 42768942 -p   R1-A_FLF_chr_allTEs_nr_noSSR_join_path -s Zt09_FLF_refTEs_seq

Concatenate .gff3 files

cat  R1-A_FLF_GFF3/*.gff3 >>  R1-A_TEs_final_annotation.gff3

Softmasking of Assembly using bedtools  (V 2.30.0)

bedtools maskfasta -soft -fi assembly.fasta -bed TEs_final_annotation.gff3 -fo assembly .softmasked_TE.fasta

Gene-Annotation using Braker 2.0 (version 2.1.6) (20) using protein homology information from the OrthoDB fungal database (Version 10).

braker.pl --cores=8 –genome= assembly.softmasked_TE.fasta --prot_seq=proteins.fasta --softmasking – fungus

Use of the Augustus Gene prediction: Augustus.gtf

Use of gffread (V0.12.7) to change to gff3 format

gffread -o augustus.hints.gff3  augustus.hints.gtf

Statistics of Proteins:  A total of 12027, 12254, and 12131 genes were predicted in the R1-A, R3-A or R3-I4 genome, respectively.

Analysis of Assembly and Annotation using BUSCO (V 5.2.2) (21).

busco -i augustus.hints.aa  -l sordariomycetes_odb10 -o busco -m protein -c 5 -f

Results of BUSCO Analysis 99.7% (R1-A: 3809/3817 BUSCO groups) or 99.8% (R3-I4: 3810/3817 BUSCO groups) or 99.8% (R3-A: 3809/3917 BUSCO groups)

The proteins were functionally annotated using Blastp Swissprot reference database using blastp (V 2.12.0):

blastp -query augustus.hints.aa -db swissprot -num_threads  -out outdir -evalue 1e-5 -outfmt 14 -taxidlist fungi.txids

For proteins encoded by genes located on chrA the number of GO Annotations were created, mapped and merged using Blast2GO (V6.0.3).

CAZymes were annotated using dbCAN2 (22) using HMMER:dbCAN with thresholds: E-Value < $1e^{-15}$, coverage >0.35) and DIAMOND: CAZy (E-Value <$1e^{-102}$) und HMMER:dbCAN-sub (E-Value <$1e^{-15}$, coverage >0,35).

### 7) Comparison with existing reads and assemblies

Publicly available whole genome sequencing reads and assemblies of 30 isolates from species of the genus Metarhizium were included in the analysis of the distribution of the accessory chromosomes within the genus. For twelve samples these were assemblies and for 18 samples these consisted of WGS reads (See Table S8 – overview of assemblies and reads, (2))

Synteny-analysis between assemblies

For the synteny analysis between assemblies the following the assemblies were aligned with nucmer (version 4.0.0rc1) and the matches were filtered for those of min 1000bp length with a minimum identity of 90%

```
nucmer --maxmatch -p outfolder R3-I4.fasta query.fasta

delta-filter -i 90 -l 1000 #.delta > #.i90.l1000.delta
```

generate bed file for coverage analysis

```
show-coords #.i90.l1000.delta > #.i90.l1000.coords

awk -v OFS='\\t' '{if ($10 >= 90 && $7 >= 90) print $12,$1-1,$2,$13-$10}'
#.i90.l1000.coords > #.i90.l1000.bed

sortBed -i #.i90.l1000.bed > #.i90.l1000.sorted.bed

bedtools merge -i  #.i90.l1000.sorted.bed > #.i90.l1000.sorted.merged.bed
```

Determining the number of SNPs in regions with coverage

```
show-snps -T -Clr  #.i90.l1000.delta > #.i90.l1000.snps

all2vcf = python mummer --snps #.i90.l1000.snps --reference  R3-I4.fasta --type SNP
--input-header --output-header > #.i90.l1000.vcf
```

For the visualization of the alignment the delta file was transformed into paf format using the paftools.js tools of minimap2 (version 2.24-r1122)

```
paftools.js delta2paf #.i90.l1000.delta  > #.i90.l1000.paf
```

Visualisation of the alignments were created using dotPlotly:

```
pafCoordsDotPlotly.R -i #.i90.l1000.paf -q 10000 -m 10000 -p 15  -s  -o out  -k 16

Rscript --vanilla pafCoordsDotPlotly.R  -i R3-I4_Mguizhouense.i90.l1000.paf -q 10000
-m 10000 -p 15 -s -o R3-I4_MguiezhouenseCologne -k 1
```

For the analysis of the coverage and SNPs of the whole genome sequencing reads deposited for 18 isolates we:

1) Deinterleaved the using the reformat.sh utility of bbmap (version 39.01)
reformat.sh in=#.fq out1=#fq_1.fastq out2=#fq_2.fastq

2) Use trimmomatic (V0.39) to remove adapter
   java -jar trimmomatic-0.39.jar PE #fq_1.fastq #fq_1.fastq #_forward_paired.fq.gz #_forward_unpaired.fq.gz #_reverse_paired.fq.gz #_reverse_unpaired.fq.gz ILLUMINACLIP:TruSeq2-PE.fa:2:30:10:2:keepBothReads LEADING:3 TRAILING:3 MINLEN:36
3) Check quality using FastQC (v0.11.5)
   fastqc #_paired.fq.gz --outdir
4) Mapping reads und sorting them using bowtie2 (version 2.4.4) and samtools (1.3.1)
```
bowtie2 -p 11 --fast -x R3-I4.fasta \
-1 #_forward_paired.fq.gz \
-2 #_forward_paired.fq.gz \
-U #_forward_unpaired.fq.gz, #_reverse_unpaired.fq.gz | samtools view -@11 -h -bS \
| samtools sort -@11 -o #-onR3-I4.sorted.bam
```
5) Some formatting and marking of duplicates of generated BAM-files using picard (version 2.18.29)
```
java -Xmx8G -jar picard.jar\
        AddOrReplaceReadGroups \
        I=#-onR3-I4.sorted.bam \
        O=#- onR3-I4.sorted.RG.bam \
        RGID=# RGLB=lib1 RGPL=ILLUMINA RGPU=unknown RGSM=#
        SORT_ORDER=coordinate"

java -Xmx8G -jar picard.jar\
MarkDuplicates \
I=#- onR3-I4.sorted.RG.bam \
O=#- onR3-I4.sorted.RG.Dedup.bam M=#_RG_Dedup.txt
```
6) Variant calling using bcftools mpileup (version= 1.14)
```
bcftools mpileup -A -f  onR3-I4.fasta #- onR3-I4.sorted.RG.bam | bcftools call --ploidy
1 -vc -Ou -o #.raw.bcf
bcftools view #.raw.bcf -o #.raw.vcf
bgzip -c #.raw.vcf > #.raw.vcf.gz && tabix -p vcf #.raw.vcf.gz
bcftools filter -o #.Q50DP6.vcf.gz  -i'QUAL>50 && DP>6' #.raw.vcf.gz
bcftools view -v snps #.Q50DP6.vcf.gz  > #.Q50DP6.snps.vcf
```

**8) SNP analysis**

Illumina reads were trimmed as described above and mapped onto the respective assemblies using bowtie2 (version 2.4.4), samtools (version 1.3.1) using the following programming:

```
bowtie2 -p 10 --sensitive -x assembly.fasta -1 paired_reads1.fq -2 paired_reads2.fq \
        -U unpaired_reads1.fq,unpaired_reads2-fq \
        |  samtools view -@10 -h -bS \
        |  sort -@10 -o out.bam
```

Formatting and marking of duplicates using PICARD (version 2.24.0)

```
java -Xmx8G -jar picard.jar \
        AddOrReplaceReadGroups \
        -INPUT out.bam \
        -OUTPUT out.RG.bam \
```

```
        -RGID sample -RGLB lib1 -RGPL ILLUMINA -RGPU unknown -RGSM sample
        -SORT_ORDER coordinate"

    java -Xmx8G -jar picard.jar \

        MarkDuplicates \

        -I out_RG.bam \

        -O out_RG_Dedup.bam -M out_RG_Dedup.txt
```

Calling of SNPs using bcftools (version 1.14)

```
    bcftools mpileup -E -C50 -Q20 -q20 -f \

        assembly.fasta out_RG_Dedup.bam | bcftools call --ploidy 1 -vc -Ou -o
        out_mpileup.raw.bcf

    bcftools view out_mpileup.raw.bcf -o out_mpileup.raw.vcf

    gzip -c out_mpileup.raw.vcf > out_mpileup.vcf.gz && tabix -p vcf
    out_mpileup.raw.vcf.gz
```

Filtering of SNPs using bcftools (version 1.14)

```
    bcftools filter -o filtered.vcf  -i 'QUAL>50 && DP>6 && AF1>0.7’ out.vcf

    bcftools filter -o indels.filtered.vcf  -i'IDV>12 && IMF>0.7' out.vcf
```

#### 9) Phasing of SNPs and small InDels for *Metarhizium guizhouense* ARSEF977

In order to phase the SNPs and small InDels of the *M. guizhouense* ARSEF977 that showed duplicated coverage of the accessory chrA *M. guizhouense* was re-sequenced PacBio HiFi chemistry. The reads were mapped onto the onR3-I4 assembly using minimap2 (version 2.24-r1122) and samtools (version 1.3.1):

```
    minimap2 -L -ax map-hifi  R3-I4.fasta  MguizhouenseARSEF977_hifi_reads.fastq.gz

    > MguizhouenseARSEF977 _on_ R3-I4.sam

    samtools view -S -b MguizhouenseARSEF977 _on_ R3-I4.sam   >

    MguizhouenseARSEF977 _on_ R3-I4.bam

    samtools sort MguizhouenseARSEF977 _on_ R3-I4.bam -o

    MguizhouenseARSEF977 _on_ R3-I4.sorted.bam
```

Phasing of SNPs and small InDels using WhatsHap (version1.6). The SNPs and small InDels called using WhatsHap (version 1.6). Please note that due to the fact that chrA of the R3-I4 assembly showed a disomic sequencing coverage the VCF the corresponding raw VCF was filtered using bcftools (version 1.14).

```
bcftools filter -o filtered.vcf  -i 'QUAL>50 && DP>10' out.vcf

whatshap phase -o Mguizhouense977_on- R3-I4_phased.vcf \
        --reference= R3-I4.fasta \
        Mguizhouense977_on- R3-I4.filtered.vcf \
        MguizhouenseARSEF977 _on_ R3-I4.sorted.bam
```

**10) Sequencing coverage analysis and distribution of SNPs in 50 kb windows**

Calculation of median coverage in 50000 bp non-overlapping windows and the fraction of the windows that is covered by at least 5 reads using mosdepth (version 0.3.3) and recovering the number of SNPs in those windows.

Generating 50 kb  windows using bedtools (version v2.25.0)

```
bedtools makewindows -b assembly.bed -w 50000  -i srcwinnum > assembly_window_50000.bed

mosdepth -t 10 -n  -T 5 -b assembly_window_50000.bed out out_RG_Dedup.bam

bedtools intersect -c -a assembly_window_50000.bed -b filtered.vcf  > SNP_50kb.counts
```

**11) Genewise relative synonymous codon usage**

Genewise relative codon usage was estimated using BioKIT (version 0.1.3) with the following command:

```
biokit gene_wise_relative_synonymous_codon_usage cds.fasta >

cds.genewise.codonusage
```

**12) Determining the Cytosine methylation within CpG context.**

To determine the whether a cytosine in a CpG context was methylated the base calling of the nanopore reads was repeated using guppy (Version 6.4.8+31becc9), minimmap2 (version 2.24-r1122) Samtools (version 1.13) was used to merge the BAM-files and mod with the model dna_r9.4.1_450bps_modbases_5mc_cg_sup.cfg.  Modified bases were called using mod_kit (version 0.1.4) with the following command:

```
modkit pileup mod.bam out.bed --cpg --ref R3-I4.fasta
```
